# DNA methylation patterns in the frontal lobe white matter of multiple system atrophy, Parkinson’s disease, and progressive supranuclear palsy: A cross-comparative investigation

**DOI:** 10.1101/2024.01.10.574852

**Authors:** Megha Murthy, Katherine Fodder, Yasuo Miki, Naiomi Rambarack, Eduardo De Pablo Fernandez, Jonathan Mill, Thomas T Warner, Tammaryn Lashley, Conceição Bettencourt

## Abstract

Multiple system atrophy (MSA) is a rare neurodegenerative disease characterized by neuronal loss and gliosis, with oligodendroglial cytoplasmic inclusions (GCI’s) containing α-synuclein being the primary pathological hallmark. Clinical presentations of MSA overlap with other parkinsonian disorders such as Parkinson’s disease (PD), dementia with Lewy bodies (DLB), and progressive supranuclear palsy (PSP), posing challenges in early diagnosis. Numerous studies have reported perturbations in DNA methylation in neurodegenerative diseases, with candidate loci being identified in various parkinsonian disorders including MSA, PD, and PSP. Although MSA and PSP present with substantial white matter pathology, alterations in white matter have also been reported in PD. However, studies comparing the DNA methylation architectures of white matter in these diseases are lacking. We therefore aimed to investigate three parkinsonian diseases, MSA, PD, and PSP, to identify shared and disease-specific DNA methylation alterations in white matter. Genome-wide DNA methylation profiling of frontal lobe white matter of individuals with MSA (n=17), PD (n=17), and PSP (n=16) and controls (n=15), using the Illumina EPIC array, revealed substantial commonalities in DNA methylation perturbations in MSA, PD, and PSP. We further used weighted gene correlation network analysis to identify disease-associated co-methylation signatures and identified dysregulation in processes relating to Wnt signalling, signal transduction, endoplasmic reticulum stress, mitochondrial processes, RNA interference, and endosomal transport. Our results highlight several shared DNA methylation perturbations and pathways indicative of converging molecular mechanisms contributing towards neurodegeneration in the white matter of all three parkinsonian diseases.

## Introduction

Multiple system atrophy is a rare adult-onset, rapidly progressing, neurodegenerative disorder characterised by neuronal loss and gliosis in multiple areas of the brain, brainstem, and spinal cord [51]. Diagnosing MSA in the early stages of the disease can be challenging owing to its overlapping clinical features with other parkinsonian disorders such as Lewy body diseases (LBD) (i.e., Parkinson’s disease (PD) and dementia with Lewy bodies (DLB)), and progressive supranuclear palsy (PSP) [34, 36, 63]. Despite sharing several neuropathological underpinnings, particularly in the advanced stages of the disease, each condition exhibits distinct neuropathological hallmarks [39]. For instance, while both MSA and PD are synucleinopathies, MSA is uniquely characterised by the presence of glial cytoplasmic inclusions (GCI’s) containing α-synuclein in oligodendrocytes, whereas PD is pathologically characterised by the presence of α-synuclein aggregates known as Lewy bodies mostly in neurons [16]. PSP, on the other hand, is a 4R tauopathy characterized by tau inclusions in the form of tufted astrocytes, neuronal tangles, and coiled bodies in oligodendrocytes, and therefore shares the common neuropathological feature of gliosis with MSA [28]. More interestingly, a co-existence of α-synuclein and tau has been observed, and both proteins share striking common characteristics suggesting a crosstalk between the two types of proteinopathies (i.e., synucleinopathies and tauopathies) and the involvement of common molecular mechanisms driving neurodegeneration [39].

The intricate mechanisms underlying neurodegeneration encompass a complex interplay between genetic, epigenetic, or regulatory factors, along with environmental exposures. Several studies have delved into the molecular underpinnings of MSA, particularly in the white matter. One such study compared grey and white matter frontal cortex transcriptomes in MSA and control subjects [37], whereas another investigated the transcriptional profiles of cerebellar white matter in MSA [50]. Epigenetic mechanisms also play a pivotal role in the tissue- and cell-type specific changes that occur during disease development and progression. DNA methylation is one of the most commonly studied epigenetic mechanisms, and perturbations in DNA methylation have been reported in several neurodegenerative disorders, including parkinsonian diseases [41]. Our group previously examined the effects of DNA methylation in white matter tissue from different brain regions in MSA compared to controls [5]. This study identified changes in key myelin and oligodendrocyte related genes, including *MOBP*, as among the most differentially methylated loci in MSA. The increased DNA methylation of *MOBP* locus observed in MSA correspondingly showed lower mRNA levels in the cerebellar white matter and although the protein levels did not differ from controls, MOBP was found to be mislocalised into the GCIs in MSA [6]. Subsequently, another study revealed a shift from cytosine methylation towards hydroxymethylation in a locus mapping to the *AREL1* gene, as well as several immune system-related changes, in the prefrontal cortex grey and white matter mixed tissue in MSA compared to controls [7, 53].

Given the clinical overlap and shared pathogenetic mechanisms between parkinsonian disorders such as MSA, PD, and PSP, a comparative analysis of DNA methylation profiles could help elucidate molecular changes common across diseases while identifying alterations specific to each pathology. To date, no study has specifically investigated DNA methylation alterations in the white matter of PD or PSP. Although PD is primarily considered to be a grey matter disease, recent transcriptomic studies have revealed an upregulation in oligodendrocyte-related genes in the brain, a downregulation in the myelin genes and the oligodendrocyte development pathway in the cingulate cortex, and a loss of oligodendrocytes in post-mortem midbrain tissue in PD patients [69]. Conversely, PSP is characterized by abnormal tau protein aggregation in both grey and white matter regions, and single-nucleus RNA sequencing in the subthalamic nucleus of PSP identified specific contributions of the glial cell-types including increased EIF2 signalling, in addition to dysregulation in genes and pathways related to apoptotic regulation and autophagy signalling in astrocytes and oligodendrocytes [18, 67]. As in the case of MSA, bulk transcriptomic analysis also revealed changes in gene expression of myelin-related genes in PSP [3]. Additionally, genetic variants in *MOBP* have been associated with PSP risk [10, 25, 54]. Oligodendrocytes are one of the major cell types in the white matter (up to 75%) that contribute to the formation of myelin sheaths and have been time and again shown to play important roles in several neurodegenerative disease mechanisms [17, 46]. Moreover, DNA methylation patterns suggest accelerated epigenetic ageing in these cells suggesting a vulnerability of oligodendrocytes to ageing [42, 43], which might, at least in part, be contributed by changes in DNA methylation.

To directly compare DNA methylation alterations in MSA white matter with those in PD and PSP, we focused on the frontal lobe, a region that is moderately affected in MSA, which also shows substantial involvement in PSP, as well as in the later stages (Braak stages 5 and 6) in PD. Additionally, previous data from our group has demonstrated a considerable overlap in DNA methylation alterations in MSA between the cerebellum (severely affected in MSA) and frontal lobe (moderately affected in MSA) [5]. Therefore, the primary objective of this study, was to perform a cross-comparative DNA methylation analysis in the frontal lobe white matter of MSA, PD, and PSP to identify distinct and shared DNA methylation alterations, and to elucidate mechanisms determining the vulnerability of specific cell types, particularly the oligodendrocytes, to dysfunction and/or protein aggregation in the different diseases.

## Material and Methods

### Human post-mortem brain tissues and their clinical and demographic characteristics

All post-mortem human brain tissues for the primary cohort were obtained from the Queen Square Brain Bank, with ethical approval for both brain donation and research protocols granted by the NRES committee – London central. The cohort comprised of human post-mortem brain tissues from individuals diagnosed with 3 neurodegenerative parkinsonian disorders, MSA (n = 17), PD (n = 17), PSP (n = 17) and neurologically healthy controls (n = 17). Disease history and neuropathological findings were characterised for all cases, and cases and controls were matched for age and sex. In addition, data from the primary cohort were compared to data previously generated by our group, which included white matter tissues from the cerebellum of individuals with MSA (n = 41) and controls (n = 21), as well as other publicly available datasets comprising grey and white matter mix tissue from the prefrontal cortex of individuals with MSA (n = 39) and controls (n = 37) (GSE143157) [53], grey matter from the frontal cortex of LBD cases (PD = 59; PD with dementia = 60; DLB = 16) and controls (n = 68) (GSE203332, GSE197305) [49], and prefrontal lobe grey and white matter from PSP cases (n = 93) and controls (n = 70) (GSE75704)[66]. Demographic and clinical characteristics for all cohorts are detailed in Table 1. GCI burden in MSA was assessed by a neuropathologist (Y.M.) using α-synuclein immunohistochemical staining in the frontal lobe white matter. The density of GCIs was graded using a modified grading scale as described previously: 0: 0-5 inclusions; 1+: 6-20 inclusions; 2+; 21-40 inclusions; 3+; ≥41 inclusions [35]. Ten areas were randomly selected in each case, and the density of GCIs was assessed using a x20 objective. Average GCI counts across the 10 areas were also used.

**Table 1:** Cohort characteristics.

| **MSA, PD and PSP frontal lobe white matter** |  |  |  |  |
| --- | --- | --- | --- | --- |
| **Group** | **Sample No.** | **Mean age (SD)** | **Sex** |  |
|  |  |  | **Female** | **Male** |
| CTRL | 17 | 72.35 (4.47) | 9 (53%) | 8 (47%) |
| MSA | 17 | 67.71 (6.47) | 8 (47%) | 9 (53%) |
| PD | 17 | 68.06 (3.67) | 9 (53%) | 8 (47%) |
| PSP | 17 | 65.35 (3.75) | 8 (47%) | 8 (50%) |
| **MSA cerebellar white matter** |  |  |  |  |
| CTRL | 21 | 80.29 (9.05) | 10 (48%) | 11 (52%) |
| MSA | 41 | 64.34 (7.9) | 20 (49%) | 21 (51%) |
| **MSA prefrontal cortex grey and white matter** |  |  |  |  |
| CTRL | 37 | 72.97 (10.45) | 18 (49%) | 19 (51%) |
| MSA | 39 | 66.03 (5.83) | 22 (57%) | 17 (43%) |
| **LBD frontal cortex grey matter** |  |  |  |  |
| CTRL | 68 | 80.41 (11.16) | 49 (72%) | 19 (28%) |
| DLB | 16 | 76.44 (8.19) | 3 (19%) | 13 (81%) |
| PD | 59 | 76.59 (8.97) | 26 (44%) | 33 (56%) |
| PDD | 60 | 78.28 (5.83) | 17 (28%) | 43 (72%) |
| **PSP prefrontal lobe grey and white matter** |  |  |  |  |
| CTRL | 70 | 76.17 (7.93) | 25 (42%) | 45 (58%) |
| PSP | 93 | 71.16 (5.32) | 39 (42%) | 54 (58%) |

CTRL – controls, MSA – multiple system atrophy, PD – Parkinson's disease, PSP – progressive supranuclear palsy, DLB – Dementia with Lewy bodies, PDD – PD with dementia

### Frontal lobe white matter DNA methylation profiling and data quality control

White matter was carefully dissected from frozen frontal lobes of individuals with MSA, PD, PSP and neurologically healthy controls and genomic DNA was extracted using a standard phenol-chloroform-isoamyl alcohol method. A total of 750 ng of DNA was subjected to bisulfite conversion using the EZ DNA Methylation kit (Zymo Research, Irvine, USA). Genome-wide DNA methylation was then performed using the Infinium HumanMethylationEPIC Bead Chip (Illumina). The resulting raw intensity (.idat) files were imported into R and subjected to thorough and stringent pre-processing and quality control checks using bioconductor packages such as minfi [4], ChAMP [57], WateRmelon [48]. Briefly, samples were evaluated by visualising raw intensities and performing outlier detection based on pcount and interquartile ranges to remove poorly performing samples; additionally, samples with a high rate of failed probes (≥2%), a mismatch in the predicted and phenotypic sex, clustering separately in the multidimensional scaling were also excluded. Probes were filtered out if they mapped to the X or Y chromosomes, were cross-reactive, of poor quality, aligned to multiple locations, or included common genetic variants. Beta-Mixture Quantile (BMIQ) normalisation method was applied to normalise the beta values, and M-values were computed as the logit transformation of beta-values, as we previously described [5].

### DNA methylation-based deconvolution of cell-type proportions

Similar to other types of genomic data, DNA methylation data derived from bulk tissue is susceptible to biases arising from variations in the cellular makeup. To address this issue, we utilized the recently developed R package ‘CEll TYpe deconvolution Goodness’ (CETYGO) [55, 62]. Building upon the functionalities of the deconvolution algorithm in the minfi package, CETYGO incorporates estimations of relative proportions of neurons (NeuN+), oligodendrocytes (SOX10+), and other glial brain cell types (Double−[NeuN−/SOX10−]) based on reference data obtained from fluorescence-activated sorted nuclei from cortical brain tissue [55]. This enabled us to estimate the cell-type proportions from frontal lobe white matter DNA methylation profiles. Comparisons between the different cell types were carried out using the Kruskal-Wallis test with a significance threshold of p-value < 0.05.

### Differential DNA methylation analysis

Given the enhanced statistical robustness of M-values [14], we employed M-values for our linear regression models using the limma package to detect differentially methylated CpG sites in MSA, PD, and PSP relative to controls, as well as between disease comparisons. To account for potential confounding factors, we incorporated age, sex, post-mortem interval (PMI), neuronal (NeuN+) proportions, and proportions of other glial cell types as covariates into the model, along with technical variables (i.e., slide, and array). The abovementioned covariates were associated with the first 5 principal components (PCs), and PCs beyond the 5^th^ PC explained <5% of the overall variance. Additionally, we utilized the SVA package [33] to estimate possible surrogate variables (SVs) and identify any unknown, latent, or unmodelled sources of noise; however, no SVs were detected using the above mentioned regression model. Adjusted beta and M-values were obtained after adjusting for the covariates included in the model described above. A false discovery rate (FDR) adjusted p-value of <0.05 was considered statistically significant at the genome-wide level, and nominal p-values ≤1 × 10^−5^ were considered suggestively significant. Adjusted beta values for CpGs that showed p < 0.0001 were used to generate heatmaps for comparisons of diseases with controls as well as comparisons between diseases.

### Weighted gene co-methylation network analysis (WGCNA)

To identify clusters of highly correlated methylation sites and to determine whether the correlation patterns were shared between the three neurodegenerative diseases, we used a systems biology method based on weighted gene correlation network analysis (WGCNA) to construct co-methylation networks [31]. The analysis utilized adjusted M-values and the top 10% CpGs mapping to genes that showed the highest variance across individuals regardless of their disease status as input (n = 53,032 CpGs). Sample clustering identified 5 outliers, which were excluded, leaving a total of 60 samples for subsequent network analysis. A signed co-methylation network was generated using the function ‘blockwiseModules’, with the ‘mergeCutHeight’ set to 0.1, soft-thresholding power of 12, and minimum module size of 200. CpGs inside each module were represented by a weighted average termed the module eigengene (ME), and highly correlated modules (ME correlation >0.75) were merged. Module membership (MM) was defined as the association between DNA methylation and each module eigengene. Additionally, we employed the applyKMeans function of the CoExpNets package [9] to reassign MM. Gene significance (GS) measure was calculated as a function GS that assigns a non-negative number to each gene; and higher GS for a gene indicates higher biological relevance or association with trait or disease. Within the disease-associated modules, we ranked genes based on their MM, prioritizing top hub genes using the function ‘chooseTopHubInEachModule’, which returns the gene with the highest connectivity in each module, looking at all genes in the methylation file [31]. Co-methylation networks were also produced in a similar way for the publicly available MSA, LBD and PSP datasets mentioned above [5, 49, 53, 66].

### Module preservation analysis in additional datasets

To assess whether the modules identified in the frontal lobe white matter were also preserved in datasets generated from different brain regions and comprising varying cell type compositions across the 3 neurodegenerative diseases, we employed preservation analysis [32]. We evaluated the module preservation for the modules identified in our data against data previously generated by our group comprising MSA cerebellar white matter, as well as with other publicly available datasets comprising MSA prefrontal cortex grey and white matter mix tissue, LBD frontal cortex grey matter tissue, and PSP prefrontal lobe grey and white matter datasets (Table 1). Preservation was determined using the ‘modulePreservation’ function of the WGCNA package with 200 permutations, and a Z-summary statistic was computed indicating high (Z-summary >10), moderate (Z-summary 2 – 10), and no preservation (Z-summary < 2) of the module in the other datasets.

### Cell type enrichment and functional network analyses for the disease-associated co-methylation modules

To delve deeper into the cellular underpinnings of the disease-associated modules, expression-weighted cell type enrichment analysis was conducted. This analysis sought to identify whether the genes within each co-methylation module were enriched for markers of a specific cell type. Leveraging the EWCE package [56] and its accompanying single-cell mouse transcriptomic dataset [70], the enrichment analysis employed p-values derived from 10,000 iterations to pinpoint enriched cell types. Subsequently, gene lists were curated for the disease-associated modules enriched for oligodendrocytes by including genes with a MM greater than 0.4, and functional module detection and enrichment analysis specific to the frontal lobe were performed using HumanBase (https://hb.flatironinstitute.org/) [22].

### Results

### Frontal lobe white matter cell-type composition across neurodegenerative parkinsonian diseases

Following our previous DNA methylation study on MSA [5], we sought to compare methylation patterns across a range of neurodegenerative parkinsonian disorders. We analysed the DNA methylation profiles generated from frontal lobe white matter from post-mortem brains of individuals with MSA, PD, PSP, and neurologically healthy controls. Following stringent quality control and filtering procedures, 65 samples (MSA = 17, PD = 17, PSP = 16, and CTRL = 15) and 734,360 probes were retained for further downstream analysis. Cell-type deconvolution methods were employed to estimate the brain cell-type proportions, confirming that the tissue samples were highly enriched for glial cells, particularly oligodendrocytes (average ∼74% across groups), consistent with the expected composition of white matter (Figure 1). Notably, in MSA and PSP, which exhibit pathological hallmarks in oligodendrocytes, we observed slightly lower proportions of oligodendrocytes, and corresponding higher proportions of other glial cells in these two diseases compared to the other groups. However, such variations in cell-type proportions failed to reach statistical significance across groups.

**Figure 1:**
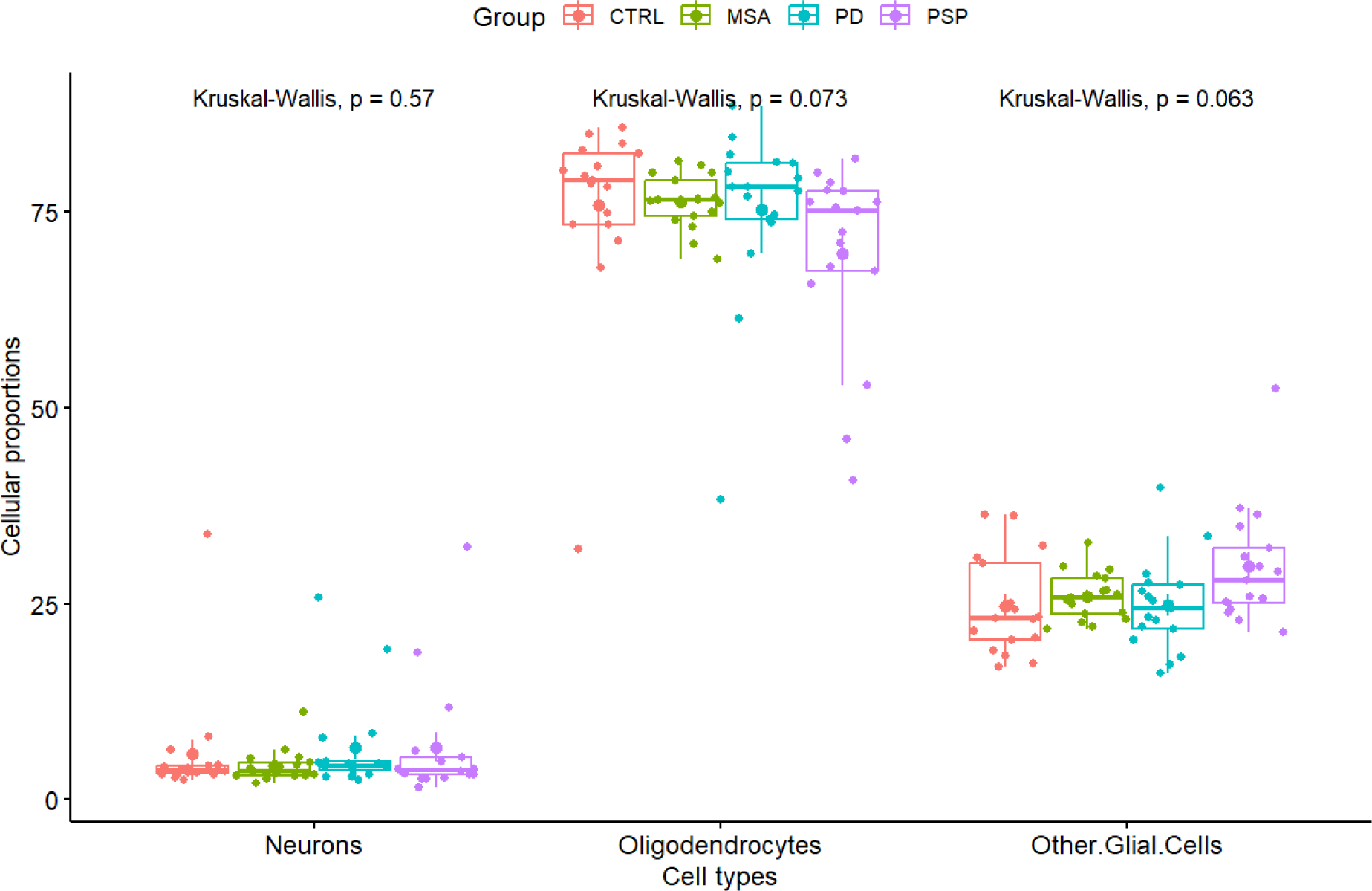
Cell-type proportion estimates for the frontal lobe white matter tissue used for DNA methylation profiling. CTRL – controls, MSA – multiple system atrophy, PD – Parkinson’s disease, PSP – progressive supranuclear palsy. Comparisons between the different sample groups were carried out using the Kruskal-Wallis test with a significance threshold of p-value < 0.05.

### Frontal lobe white matter DNA methylation profiling shows shared patterns across neurodegenerative parkinsonian diseases

We next conducted an epigenome-wide association study (EWAS) using a linear regression model that accounted for age at death, sex, cell proportions, and other covariates as detailed in the methods. Quantile-quantile plots showed no evidence of genomic inflation for any of the comparisons (Figure S1). When considering the topmost differentially methylated sites (p<0.0001) in all three parkinsonian diseases (MSA, PD, and PSP together) versus controls (Table S1), a clear separation was observed between parkinsonian diseases and controls (Figure 2a). However, little or no separation was observed within the three disease groups. Although not passing genome-wide significance after multiple testing corrections, eight CpGs mapping to seven genes (*SFI1*, *IL22RA2*, *WWOX*, *ETNK1*, *CEP41*, and *FAM8A1*, *C4orf50*) showed shared differential methylation (hypo- or hypermethylation) with a suggestive significance of p≤1 × 10^−5^ across all diseases and interestingly, several of these genes have been previously associated with neurological conditions (Figure 2b,c, Table 2).

**Figure 2:**
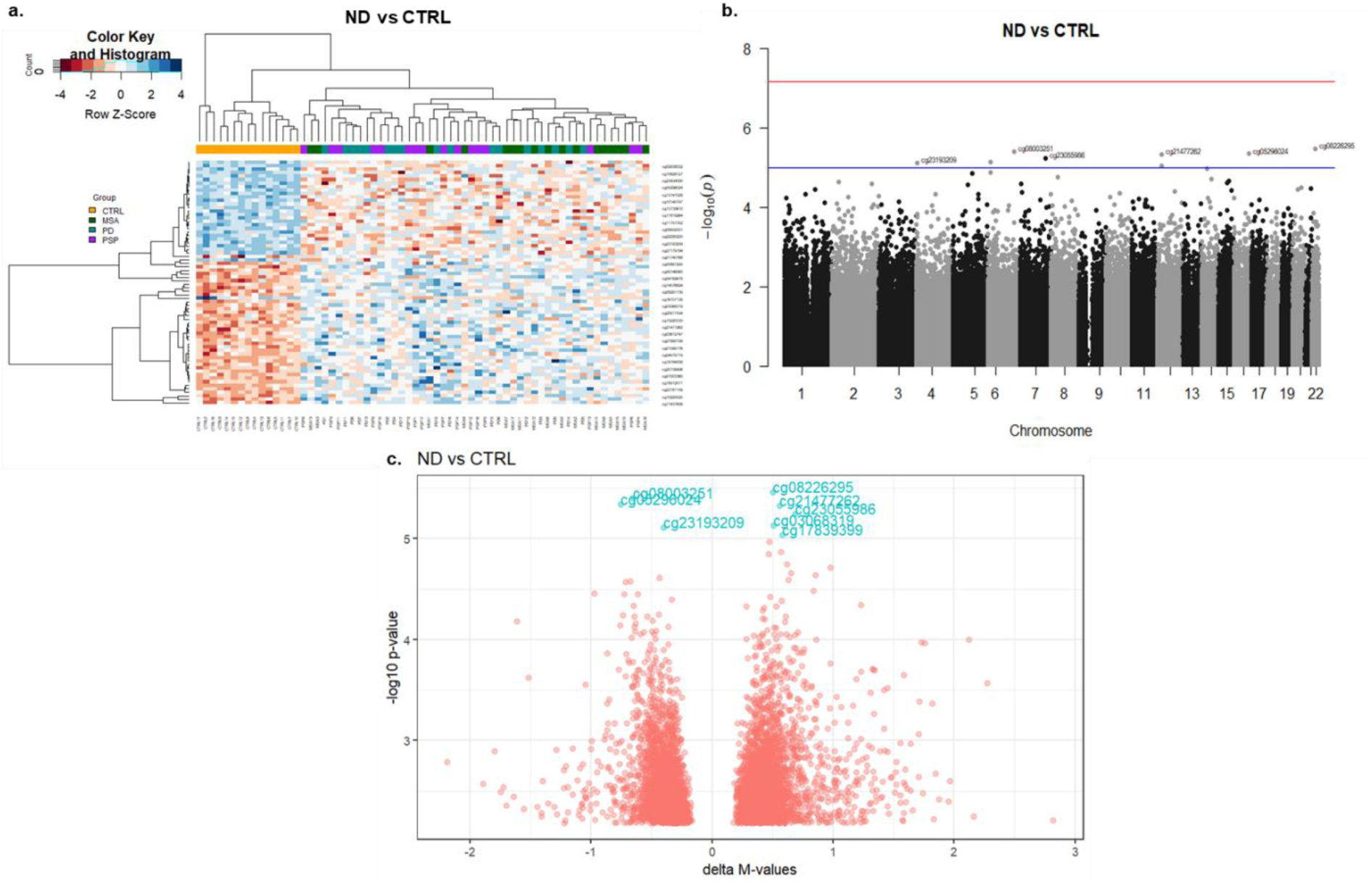
DNA methylation alterations in the frontal lobe white matter shared across the three neurodegenerative parkinsonian diseases (MSA, PD, and PSP). (a) Heatmap of the topmost significant differentially methylated loci (p < 0.0001) identified in the neurodegenerative parkinsonian diseases compared to controls. The rows represent CpGs, columns represent samples, and the colours represent the direction as well as the magnitude of effect (adjusted β values) in all the samples (darker colours indicate larger effect sizes). (b) Manhattan plot showing associations between single DNA methylation sites (CpGs) and the neurodegenerative parkinsonian diseases. The red line indicates genome-wide significance threshold based on Bonferroni-corrected p-values (p = 6.8 × 10^−8^), and the blue line indicates a less stringent suggestive significance threshold of p = 1 × 10^−5^. (c) Volcano plot showing the differentially methylated probes shared across the neurodegenerative parkinsonian diseases, CpGs with suggestive significance threshold of p = 1 × 10^−5^ are highlighted in blue. CTRL – controls, ND – Neurodegenerative parkinsonian diseases.

**Table 2:** Differentially methylation positions (DMPs) identified with suggestive significance (p<1 × 10^−5^) in the different group comparisons.

| CpGs | Gene Symbol | Delta Beta (Adj) | Delta M-val | P.Value | adj.P. Val | Chr | Position | Feature | cgi |
| --- | --- | --- | --- | --- | --- | --- | --- | --- | --- |
| ND vs CTRL |  |  |  |  |  |  |  |  |  |
| cg08226295 | *SFI1* | 0.003 | 0.50 | 3.44E-06 | 0.82 | 22 | 31892562 | 5'UTR | island |
| cg08003251 | *IL22RA2* | -0.02 | -0.65 | 3.99E-06 | 0.82 | 6 | 137469991 | Body | opensea |
| cg05296024 | *WWOX* | -0.002 | -0.75 | 4.56E-06 | 0.82 | 16 | 78495308 | Body | opensea |
| cg21477262 | *ETNK1* | 0.01 | 0.56 | 4.68E-06 | 0.82 | 12 | 22777430 | TSS1500 | shore |
| cg23055986 | *CEP41* | 0.01 | 0.69 | 5.72E-06 | 0.82 | 7 | 130080444 | Body | shore |
| cg03068319 | *FAM8A1* | 0.02 | 0.51 | 7.37E-06 | 0.82 | 6 | 17600252 | TSS1500 | shore |
| cg23193209 | *C4orf50* | -0.01 | -0.40 | 7.78E-06 | 0.82 | 4 | 5972071 | Body | opensea |
| cg17839399 | *ETNK1* | 0.01 | 0.58 | 9.14E-06 | 0.84 | 12 | 22777409 | TSS1500 | shore |
| **MSA vs CTRL** |  |  |  |  |  |  |  |  |  |
| cg22290225 | *MEGF11* | -0.02 | -0.62 | 1.08E-06 | 0.80 | 15 | 66319931 | Body | opensea |
| **cg15274294** |  | -0.07 | **-0.80** | **5.12E-06** | **0.94** | **6** | **14884128** | **IGR** | **opensea** |
| **cg15644686** | ***BCL7B*** | -0.33 | **-2.19** | **9.52E-06** | **0.94** | **7** | **72972216** | **TSS1500** | **island** |
| **PD vs CTRL** |  |  |  |  |  |  |  |  |  |
| cg10828127 | *DSCAM* | -0.003 | -1.40 | 4.15E-07 | 0.30 | 21 | 41550814 | Body | shelf |
| **cg01380065** | ***UBE2F*** | **0.10** | **1.33** | **1.25E-06** | **0.46** | **2** | **238878242** | **5'UTR** | **shore** |
| cg11757352 |  | -0.01 | -0.64 | 4.63E-06 | 0.89 | 13 | 30728812 | IGR | opensea |
| cg03068319 | *FAM8A1* | 0.02 | 0.56 | 9.88E-06 | 0.89 | 6 | 17600252 | TSS1500 | shore |
| **PSP vs CTRL** |  |  |  |  |  |  |  |  |  |
| cg04596067 | *MYT1L* | -0.003 | -0.81 | 4.34E-06 | 0.99 | 2 | 2026609 | 5'UTR | opensea |
| cg04956571 | *CBX8* | 0.001 | 0.79 | 5.28E-06 | 0.99 | 17 | 77770388 | Body | shore |
| cg17839399 | *ETNK1* | 0.02 | 0.75 | 5.37E-06 | 0.99 | 12 | 22777409 | TSS1500 | shore |
| cg23193209 | *C4orf50* | -0.02 | -0.52 | 5.69E-06 | 0.99 | 4 | 5972071 | Body | opensea |
| cg12824502 | *MPI* | 0.00 | 0.89 | 6.85E-06 | 0.99 | 15 | 75182590 | Body | island |
| **cg25358066** | ***D2HGDH*** | 0.07 | **0.59** | **9.30E-06** | **0.99** | **2** | **242695249** | **ExonBnd** | **opensea** |
| cg10377240 | *FAM179A* | -0.01 | -0.83 | 9.41E-06 | 0.99 | 2 | 29248433 | Body | opensea |
| **MSA vs PD** |  |  |  |  |  |  |  |  |  |
| **cg05376227** | ***FMO6P*** | **0.08** | **0.70** | **1.85E-06** | **0.74** | **1** | **171111193** | **Body** | **opensea** |
| cg07377662 | *METRNL* | 0.01 | 0.42 | 2.24E-06 | 0.74 | 17 | 81037199 | TSS1500 | island |
| cg24624576 | *SEC63* | 0.01 | 0.65 | 4.36E-06 | 0.74 | 6 | 108224767 | Body | opensea |
| cg12055395 | *DLX6AS* | 0.02 | 1.09 | 4.86E-06 | 0.74 | 7 | 96642605 | Body | shelf |
| cg07840454 | *VN1R1* | 0.04 | 0.57 | 6.85E-06 | 0.74 | 19 | 57968785 | TSS1500 | opensea |
| cg02434357 | *SERINC4* | -0.002 | -0.77 | 7.41E-06 | 0.74 | 15 | 44093254 | TSS1500 | shore |
| cg14932313 | *CTSG* | 0.03 | 0.46 | 8.63E-06 | 0.74 | 14 | 25043420 | Body | opensea |
| **cg20311843** | ***OR51A7*** | **0.07** | **0.61** | **9.90E-06** | **0.74** | **11** | **4927620** | **TSS1500** | **opensea** |
| **MSA vs PSP** |  |  |  |  |  |  |  |  |  |
| cg04956571 | *CBX8* | -0.001 | -0.80 | 1.31E-06 | 0.96 | 17 | 77770388 | Body | shore |
| cg22713693 | *CUBN* | 0.01 | 0.54 | 3.17E-06 | 1.00 | 10 | 16895852 | Body | opensea |
| cg02434357 | *SERINC4* | -0.003 | -0.92 | 7.29E-06 | 1.00 | 15 | 44093254 | TSS1500 | shore |
| **PD vs PSP** |  |  |  |  |  |  |  |  |  |
| **cg06831571** |  | **0.12** | **0.90** | **3.82E-06** | **1.00** | **11** | **34592196** | **IGR** | **opensea** |
| cg00843912 | *ZNF180* | -0.02 | -0.76 | 4.64E-06 | 1.00 | 19 | 45004550 | 1stExon | island |
| cg07366967 | *EXD2* | 0.03 | 0.63 | 7.47E-06 | 1.00 | 14 | 69675438 | TSS1500 | opensea |

ND – Neurodegenerative parkinsonian diseases, CTRL – controls, MSA – multiple system atrophy, PD – Parkinson’s disease, PSP – progressive supranuclear palsy. CpGs highlighted in bold show an absolute delta beta values ≥5%.

### Specificities of frontal lobe white matter DNA methylation profiles in each of the neurodegenerative parkinsonian disorders

To identify the DNA methylation changes of higher relevance to each disease, we also compared the individual disease groups with controls and identified the top most-differentially methylated CpGs (p≤1 × 10^−5^), which included 3 CpGs in MSA, 4 CpGs in PD, and 7 CpGs in PSP (Figure S2a-c, Table 1). Among these differentially methylated CpGs, only cg15274294, and cg15644686 (*BCL7B*) in MSA, cg01380065 (*UBE2F*) in PD, and cg25358066 (*D2HGDH*) in PSP, showed substantial effect sizes with absolute delta beta values ≥5% (Figure 3); however, the direction of effect for these CpGs remained the same in all three diseases. Notably, there was some overlap of the CpGs identified in the individual disease groups with the CpGs identified in the overall comparison, including CpGs in *FAM8A1*, *C4orf50*, and *ETNK1*. Interestingly, in line with the expected downstream effect of hypomethylation the promoter region of *BCL7B* (cg15644686) on gene expression levels, a previous study on the transcriptional profiling of cerebellar white matter in MSA reported an upregulation of *BCL7B* (log2FC = 0.576, adj.P= 2.4 × 10^−2^) [50].

**Figure 3:**
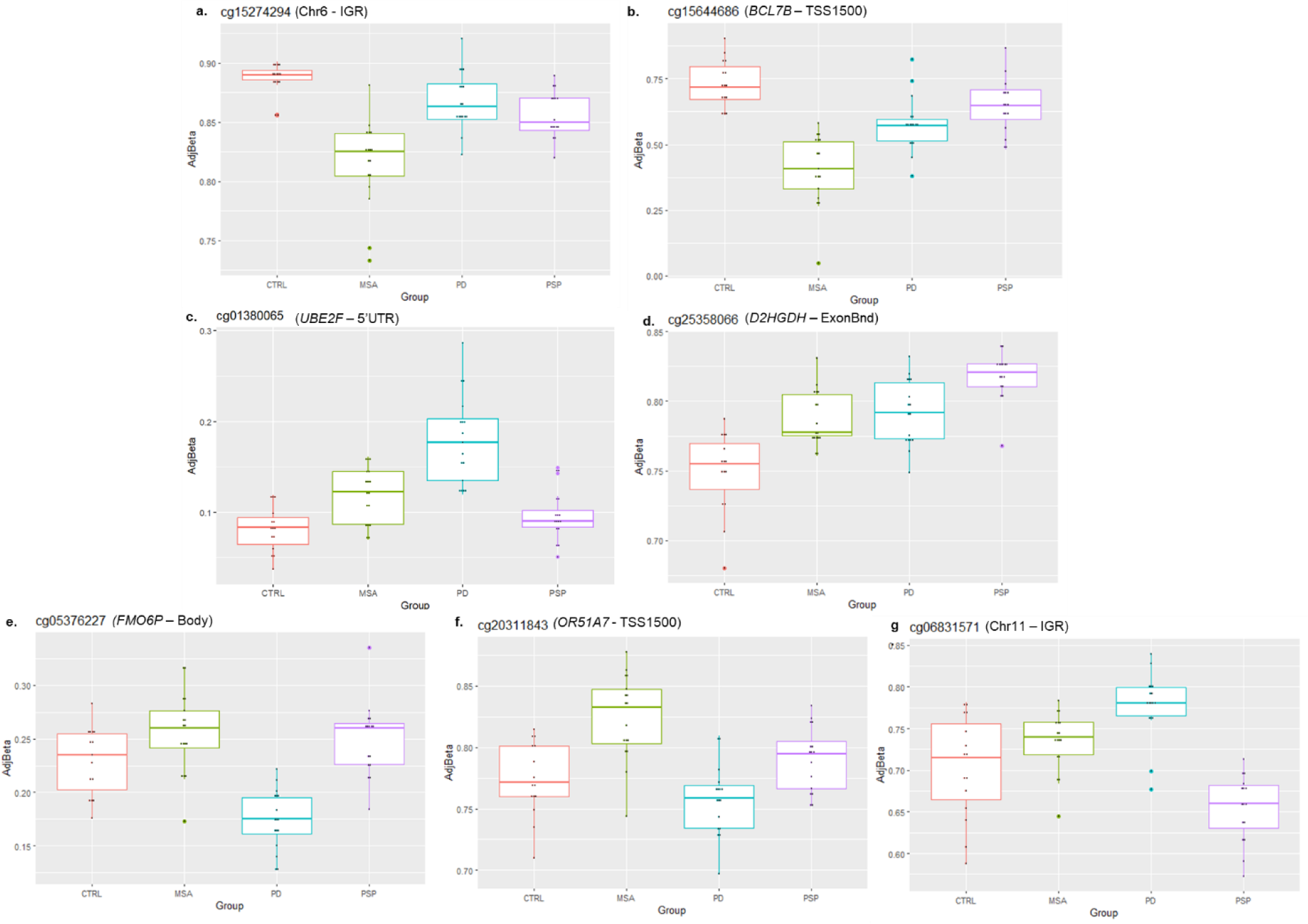
DNA methylation levels for the differentially methylated positions (DMPs) in neurodegenerative parkinsonian diseases versus controls showing suggestive significance (p<1 × 10^−5^) and effect size (absolute delta beta values) ≥ 5%; (a, b) hypomethylation at cg15274294 (Chr 6 – IGR), and cg15644686 (*BCL7B*) in MSA vs controls, (c) hypermethylation at cg01380065 (*UBE2F*) in PD vs controls, (d) hypermethylation at cg25358066 (*D2HGDH*) in PSP vs controls, (e, f) hypermethylation at cg05376227 (*FMO6P*), cg20311843 (*OR51A7*) in MSA vs PD and (g) hypermethylation at cg06831571 (Chr11 – IGR) in PD vs PSP. CTRL – control, MSA– multiple system atrophy, PD – Parkinson’s disease, PSP – progressive supranuclear palsy, IGR – intergenic region, TSS1500 – 200-1500 bases upstream of the transcription start site, ExonBnd – exon boundary.

As the majority of the differentially methylated CpGs identified within the disease groups compared to controls showed a similar direction of effect in the three disease groups (Figure S3a–c), we also compared the disease groups against each other to investigate whether there are differential methylation signatures with potential to discriminate between the disease groups (MSA vs PD, MSA vs PSP, and PD vs PSP). Among the top most-differentially methylated CpGs in the three parkinsonian diseases (Table 1, Figure S2d-f), only cg05376227 (*FMO6P*), and cg20311843 (*OR51A7*) in MSA vs PD, and cg06831571 (Chr11 – IGR) in PD vs PSP showed effect sizes with absolute delta beta values ≥5% (Figure 3). However, for these CpGs, only the two disease groups being compared showed opposite direction of effects, whereas the other two were always concordant. The overall analysis of the top ranked differentially methylated positions (p < 0.0001) for each comparison also revealed more similarities in DNA methylation patterns between MSA and PD (Figure S3d), both synucleinopathies, compared to that between MSA and PD vs PSP (Figure S3e, f), which is a tauopathy.

### Top differentially methylated positions in parkinsonian diseases are associated with disease traits

We also explored the top differentially methylated positions (DMPs) with effect size >5% between each disease and controls further to identify associations with disease traits such as the average number of glial cytoplasmic inclusions (GCIs) in the oligodendrocytes in case of MSA, as well as disease onset and duration for all diseases. We found that methylation levels at the intergenic cg15274294 in MSA was inversely associated with the mean number of GCIs in the frontal lobe (R=-0.58, p=0.029) (Figure S4a), and the same direction of effect was observed for disease duration (R=-0.34, n.s.); in PD, lower methylation levels in this CpG also showed significant correlations with earlier onset of disease (R=0.59, p=0.013), but longer disease duration (R=-0.50, p=0.042) (Figure S4b,c). Overall, these findings suggest that the methylation status at this site is related with disease progression in synucleinopathies. For cg15644686 (*BCL7B*), which showed the strongest biological effect with MSA status (Figure S4a), despite not reaching statistical significance, an inverse relationship was observed between the methylation levels and the mean number of GCIs (R=-0.36, n.s.) as well as with the disease duration (R=-0.27, n.s.). For cg01380065 (*UBE2F*), which showed the strongest biological effect with PD status, higher methylation levels tended to be associated with delayed disease onset (R=0.32, n.s.) and shorter disease duration (R=-0.36, n.s.) (Figure S4b,c). For cg25358066 (*D2HGDH*), which showed the strongest biological effect with the PSP status, higher methylation levels also tended to be associated with a shorter disease duration in PSP (R=-0.47, n.s.) (Figure S4c).

Among the CpGs showing the strongest effects in the comparisons within disease groups, a significant negative correlation was also observed between methylation levels in cg05376227 (*FMO6P*) and avgGCI in MSA (Figure S5a); however, no significant correlations were observed with disease onset or duration (Figure S5b,c).

### WGCNA identifies shared and disease specific DNA co-methylation modules

We further performed co-methylation analysis using WGCNA to identify clusters of highly correlated CpGs (modules) (Figure S6). A total of 32 co-methylation modules were identified, of which 15 modules were significantly associated with the status of at least one disease group (p≤0.0015, 0.05/32 modules) (Figure 4a, Figure S7a). Among these, the lightcyan module was positively associated with all three disease groups [MSA (R=0.71, p=2 × 10^−10^), PD (R= 0.67, p=6 × 10^−9^), and PSP (R=0.75, p=8 × 10^−12^)] (Figure 4b), whereas the darkgrey module showed a positive correlation in both PD and PSP [PD (R=0.43, p=6 × 10^−4^); PSP (R=0.66, p=1 × 10^−8^)] and to a certain extent in MSA (R=0.32, p=0.01). Modules significantly associated only with α-synucleinopathies included darkturquoise [MSA (R=0.56, p=3 × 10^−6^); PD (R=0.51, p=3 × 10^−5^)], darkgreen [MSA (R=-0.57, p=3 × 10^−6^); PD (R=-0.45, p=3 × 10^−4^)], and white [MSA (R=-0.62, p=1 × 10^−7^); PD (R=-0.61, p=3 × 10^−7^)] (Figure 4c). As with the differential methylation analysis, both synucleinopathies had more similarities among them than with PSP, with concordant direction of effects in all disease associated modules for MSA and PD (Figure 4).

**Figure 4:**
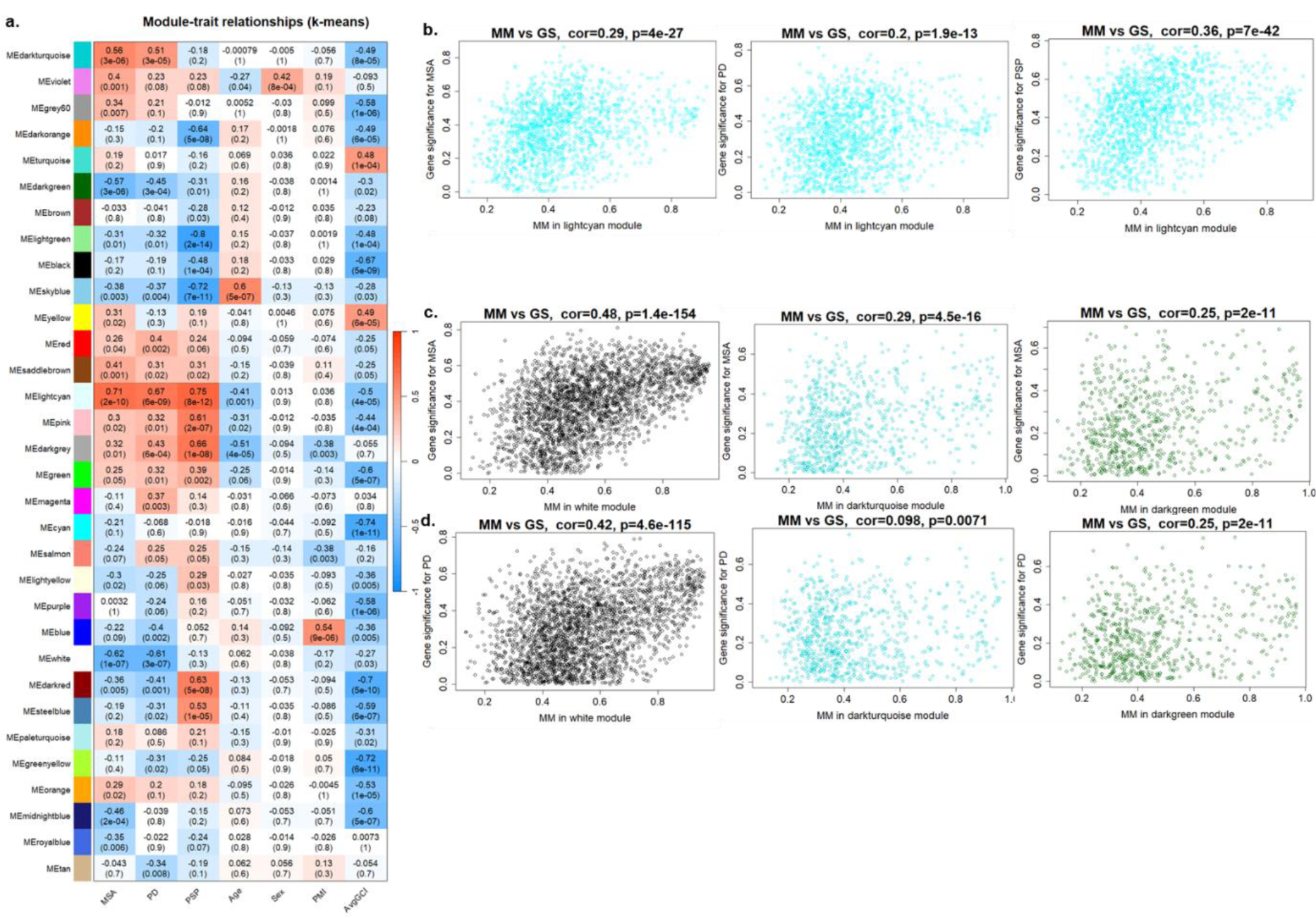
Co-methylation network analysis. (a) module-trait correlations for the co-methylation networks. Rows represent co-methylation module eigengenes and their colours; columns represent the correlation (and p-values) of the methylation levels of CpGs in each module with the disease status and clinical/pathological traits. Colour scale at the right indicates the strength of the correlation (darker cells depict stronger correlations, with blue representing negative and red representing positive correlations). Gene significance versus module membership (kME) for (b) lightcyan module which was significantly associated (adj.p≤0.001) with all three disease groups (MSA, PD, PSP), (c,d) modules significantly associated (adj.p≤0.001) with α-synucleinopathies (MSA and PD) (white, darkturquoise, and darkgreen). MSA – multiple system atrophy, PD – Parkinson’s disease, PSP – progressive supranuclear palsy, GS – gene significance, MM – module membership.

Modules significantly associated with MSA only included violet (R=0.4, p=0.001), saddlebrown (R=0.41, p=0.001) among the positively correlated modules, and midnightblue (R=-0.46, p=2 × 10^−4^) among the negatively correlated ones (Figure 4, Figure S8a). The darkred module was negatively associated with PSP and showed an inverse association with both α-synucleinopathies [MSA (R=-0.36, p=0.005), PD (R=-0.41, p=0.001); PSP (R=0.63, p=5 × 10^−8^)]. The steelblue module (R=0.53, p=1 × 10^−5^) was positively associated with PSP only (R=0.53, p=1 × 10^−5^) but showed non-significant inverse correlations with both α-synucleinopathies [MSA (R=-0.19, n.s.), PD (R=-0.31, p=n.s.)] (Figure 4, Figure S8a-c). Other modules significantly associated with PSP, but showing similar non-significant associations with MSA and PD included the positively correlated pink module (R=0.61, p=2 × 10^−7^), and negatively correlated modules lightgreen (R=-0.8, p=2 × 10^14^), skyblue (R=-0.72, p=7 × 10^−11^), darkorange (R=-0.64, p=5 × 10^−8^), and black (R=-0.48, p=1 × 10^−4^) (Figure 4, Figure S8a-c). No PD specific modules were identified.

We also investigated whether the disease-associated modules were associated with clinical and pathological traits such as age of disease onset and disease duration, and the average number of GCIs (avgGCI) in MSA. The darkred module positively associated with PSP also showed negative correlation with disease duration in the PSP samples suggesting that higher methylation levels in the CpG sites within this module could contribute towards a faster progression and shorter disease duration (Figure S7a). Among the modules significantly associated with the MSA status, the darkturquoise, the lightcyan, and the midnightblue also showed negative correlations with avgGCI (Figure 4), supporting a role of DNA methylation in disease progression.

We further explored each of the modules significantly associated with one or more disease groups and investigated module memberships to identify intramodular hub genes. Hub gene analysis further identified dysregulation in several genes previously implicated in neurodegeneration as well as in parkinsonian diseases (Table S2). Interestingly, the pyroptotic gene *DFNA5* was identified as the top hub gene in the lightcyan module commonly dysregulated in the three diseases. Hub genes associated with a-synucleinopathy associated modules included *RBP4, C1orf70 (TMEM240),* and *SCARF2,* which have been previously implicated in PD [65].

### Disease associated co-methylation modules enriched for oligodendrocytic gene signatures display enrichment of distinct molecular pathways involved in neurodegeneration

As we analysed DNA methylation in the white matter, we performed cell-type enrichment analysis to identify modules significantly enriched for specific glial cells and further understand their contribution disease related processes. The darkgrey, darkred, steelblue, and white modules showed a significant enrichment in oligodendrocytic gene signatures (Figure 5a). When assessing enrichment for specific subpopulations within the oligodendrocyte lineage, the midnightblue module was specifically enriched for the sub-cell type Oligo1, which is thought to correspond to oligodendrocytes undergoing differentiation (Figure 5b) [70]. The steelblue and white modules showed significant enrichments for Oligo2 and Olig6, which represent pre-myelinating and terminally differentiated post-myelinating oligodendrocytes, respectively. No significant enrichment was observed for astroglial and microglial cell types. Neuronal proportion estimates within the dataset were negligible, and therefore, any enrichment for neuronal markers was interpreted as related with neuronal signatures being silenced in the white matter.

**Figure 5:**
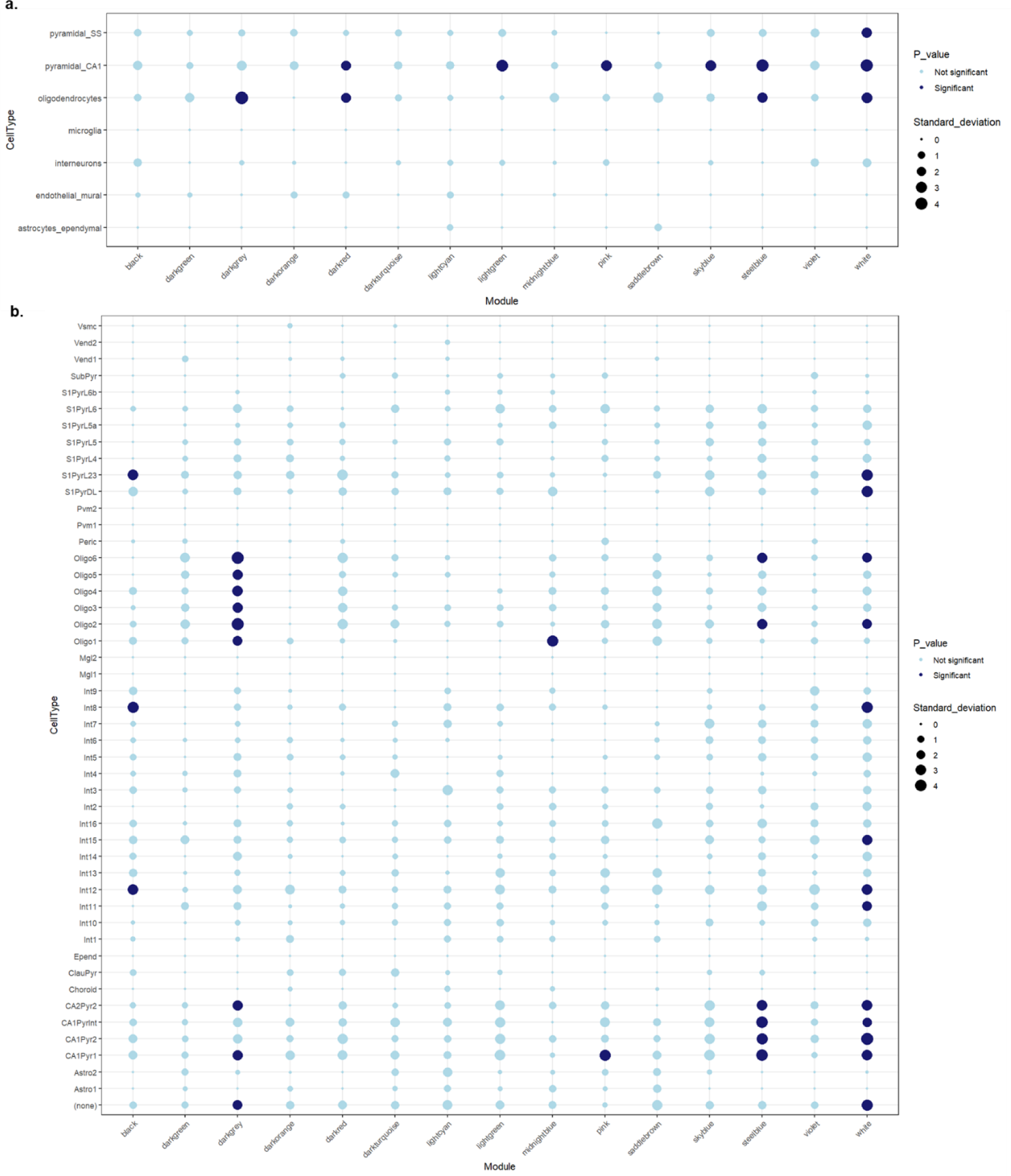
Cell-type enrichment for the WGCNA modules associated with one or more disease groups. Enrichment for the different brain (a) cell types and (b) cell subtypes performed using the package EWCE and associated single-cell transcriptomic data which uses mouse to human homologs of genes associated with various cell types; dark blue circles represent significantly enriched cell types with adjusted p < 0.05 after Bonferroni corrections; the size of the circles represents the number of standard deviations (SD) from the mean.

As our main objective was to elucidate the contribution of DNA methylation perturbation to the parkinsonian diseases in tissues enriched for specific glial cell types, we focused our attention on further exploring modules specifically associated with oligodendrocyte signatures. We therefore performed gene ontology enrichment and functional network analysis specific for the frontal lobe for genes in the oligodendrocyte enriched co-methylation modules to understand their functional significance. The darkgrey module, which was significantly associated with PD and PSP and also to a certain extent with MSA showed submodules enriched for processes such as RNA interference (M1), immune response-activating signal transduction (M4), regulation of exosomal secretion, response to endoplasmic reticulum (ER) stress, regulation of mitochondrial translation (M5), and endosomal transport (M6) (Figure 6a, Table S3). The white module significantly associated with only the α-synucleinopathies (MSA and PD) showed its largest submodule (M3) to be enriched for processes such as regulation of Wnt signalling pathway and cell-cell signalling by Wnt (Figure 6b, Table S3). Interestingly, this submodule had *PARKIN* (a causal gene in familial forms of PD [11]) as its hub gene, and also contained the gene *BCL7B,* to which our top DMP cg15644686 in MSA mapped. The darkred and the steelblue modules showed negative associations with α-synucleinopathies and positive association with PSP. The largest submodule in darkred (M8) was enriched for processes involved with histone methylation, cell migration, and Wnt signalling pathway, again with *PARKIN* as its hub gene (Figure 6c, Table S3). The steelblue module was enriched for processes such as protein localisation to nucleoplasm (M2), RNA splicing and translation (M1), and antigen processing and presentation (M3) (Figure 6d, Table S3). The midnightblue module was the only module exclusively associated with MSA and showed enrichment for negative regulation of Wnt signalling pathway, cellular response to lipid, and regulation of SMAD protein signal transduction in the M4 submodule and processes such as regulation of protein dephosphorylation, protein targeting and localisation to mitochondrion and apoptotic signalling in response to mitochondrial stress (Figure 6e, Table S3).

**Figure 6:**
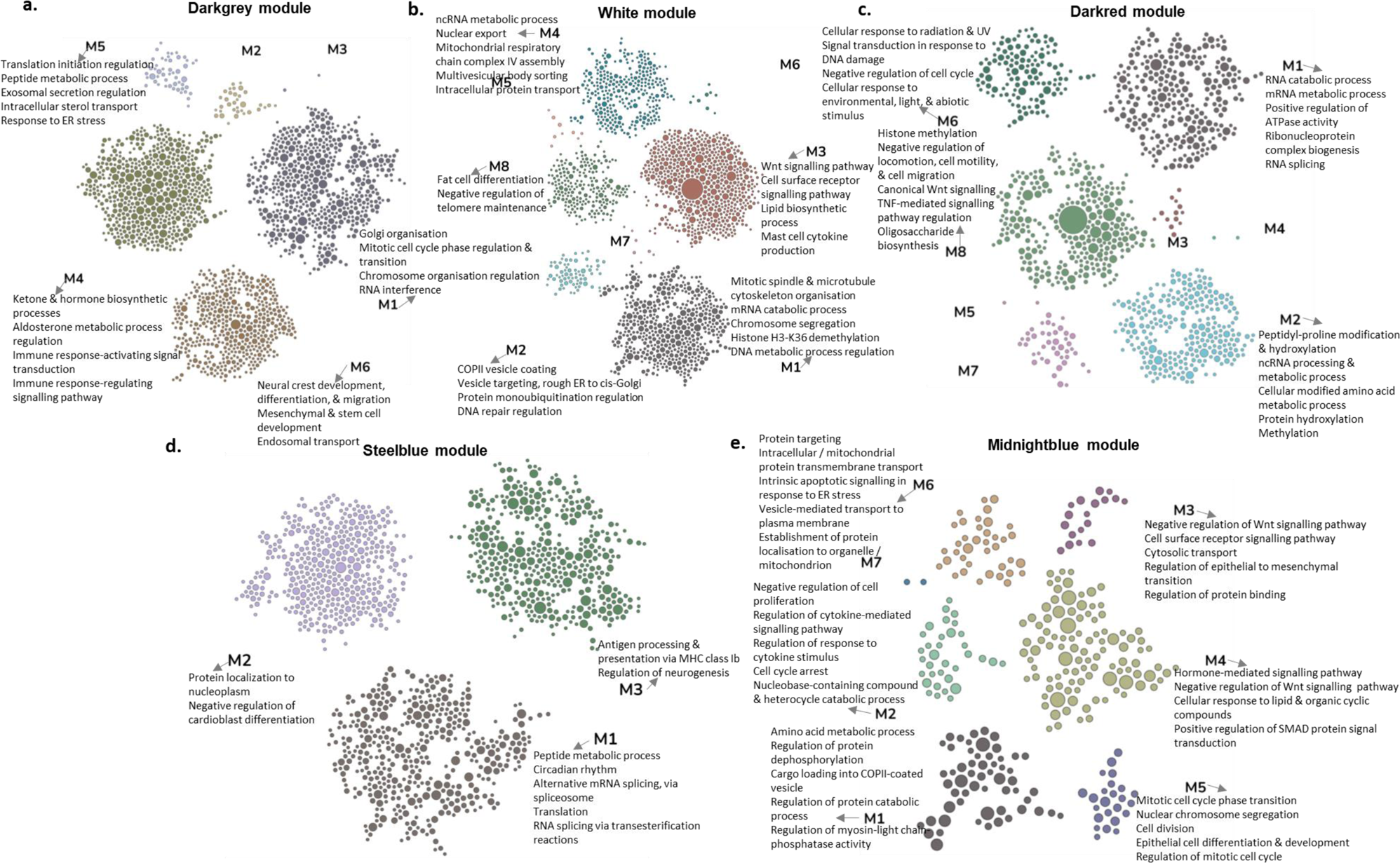
Frontal lobe specific functional network analysis on the oligodendrocyte enriched modules created using HumanBase (https://hb.flatironinstitute.org/) for the (a) darkgrey module associated with all three disease groups (MSA, PD, PSP), (b) white module significantly associated with α-synucleinopathies (MSA and PD), (c) darkred and (d) steelblue modules negatively associated with α-synucleinopathies (MSA and PD) but positively associated with PSP, and (e) midnightblue module exclusively associated with MSA. Enrichment for top 5 relevant terms within the functional modules of each WGCNA modules are listed. MSA– multiple system atrophy, PD – Parkinson’s disease, PSP – progressive supranuclear palsy.

### Disease-associated modules are preserved to various degrees in other brain regions and tissue types in MSA, Lewy body diseases (LBD), and PSP, and display overlaps in enriched processes

To assess replication of parkinsonian disease-associated white matter co-methylation modules, we performed preservation analysis using multiple previously available datasets from other brain regions, tissue types, and diseases. We compared our data with DNA methylation profiles generated previously by our group for MSA for cerebellar white matter, and publicly available datasets of MSA prefrontal cortex grey and white matter, LBD frontal cortex grey matter, and PSP prefrontal lobe grey and white matter datasets. Most of the MSA associated modules (4-5/7) showed moderate preservation (Z-summary > 2) in the other two MSA datasets, including the oligodendrocyte associated midnightblue and white modules in MSA for cerebellar white matter and white module in MSA prefrontal cortex grey and white matter (Figure 7a,b). All the PD associated modules showed high or moderate preservation in the LBD frontal cortex grey matter dataset, with the oligodendrocyte enriched modules darkgrey, darkred, and white being highly preserved (Z-summary > 10) (Figure 7c). Most modules associated with PSP (6/9) also displayed moderate preservation in the PSP prefrontal lobe grey and white matter dataset and included oligodendrocyte enriched modules darkred and steelblue (Figure 7d).

**Figure 7:**
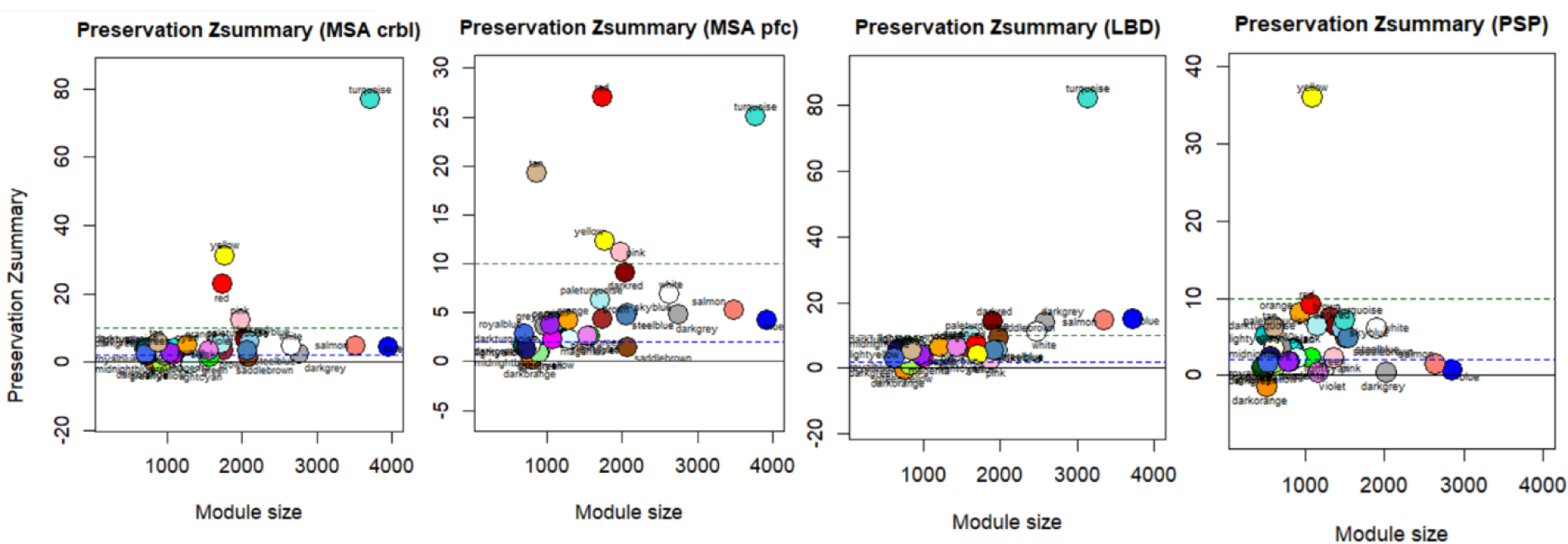
Preservation analysis for the co-methylation modules in the frontal lobe white matter dataset for MSA, PD, and PSP in other brain regions and tissue types. Preservation Z summaries of the co-methylated modules in (a) MSA cerebellar white matter dataset, (b) MSA prefrontal cortex grey and white matter dataset, (c) Lewy body disease (LBD) frontal cortex grey matter dataset, and (d) PSP prefrontal lobe grey and white matter dataset. Y-axis represents the preservation Z-summary with modules above the green dashed line (Z-summary >10) predicted to be highly preserved, those between the blue and green dashed lines (2< Z-summary <10) predicted to be moderately preserved, and modules below the blue line (Z-summary <2) are not preserved. MSA– multiple system atrophy, PD – Parkinson’s disease, PSP – progressive supranuclear palsy.

We further performed WGCNA on the abovementioned additional datasets to identify functional similarities between disease-associated modules identified in our dataset and those identified in the other datasets comprising different brain regions and tissue-types. Several overlapping disrupted processes and pathways were identified from the functional enrichment and network analyses of these datasets (Table S4). Modules associated with MSA in the cerebellar white matter dataset and MSA in our dataset demonstrated shared enrichment for processes such as protein phosphorylation, cell migration, and cell motility. In case of the MSA prefrontal cortex grey and white matter mix dataset, common enriched pathways included protein dephosphorylation, regulation of peroxisome organization, and intracellular protein transport, among others. Modules associated with LBD frontal cortex grey matter and PD in our dataset exhibited shared processes such as transmembrane receptor protein tyrosine kinase signalling pathway. Enriched processes common between modules associated with PSP prefrontal cortex grey and white matter and PSP in our dataset included regulation of mRNA metabolic process, DNA metabolic process, and chromosome organization, response to endoplasmic reticulum stress, histone methylation, endosomal transport, and positive regulation of TOR signalling (Table S4).

## Discussion

We performed a cross-comparative analysis of DNA methylation changes in the frontal lobe white matter of individuals with MSA, PD, and PSP to identify shared and disease-specific molecular signatures in the white matter. Despite the variable extent of white matter involvement across these three parkinsonian diseases [12], a comprehensive analysis revealed substantial commonalities in DNA methylation alterations, with a majority of the differentially methylated CpGs displaying a similar direction of effect, albeit with varying effect sizes. This shared DNA methylation architecture suggests that the presence of similar pathogenic mechanisms and cellular responses in MSA, PD, and PSP extends to (cell-types within) the white matter. Among the three parkinsonian diseases, MSA and PD exhibited greater similarities compared to MSA and PSP or PD and PSP, suggesting that the synucleinopathies share more commonalities, despite differences in the cell-types where α-synuclein inclusions primarily occur. Additionally, this observation suggests that despite the more extensive white matter involvement in MSA and PSP relative to PD [45], the aggregated protein type (α-synuclein vs. tau) introduces a greater degree of difference in the underlying molecular processes, as reflected in the DNA methylation patterns across the three diseases.

The top differentially methylated CpGs identified with suggestive significance commonly altered in MSA, PD, and PSP, mapped to several genes with prior associations with neurological conditions. For instance, *IL22RA2,* a multiple sclerosis risk gene, encodes the soluble interleukin (IL)-22-binding protein (IL-22BP) [8] and higher levels of IL-22 in multiple sclerosis have been shown boost Fas expression in oligodendrocytes, resulting in oligodendrocytic apoptosis [71]. *WWOX*, which is a risk gene for AD and also implicated in PD and multiple sclerosis [2, 26], has been shown to modulate GSK3β, EKR, and JNK kinase activities responsible for Tau hyperphosphorylation resulting in aggregation of Tau into neurofibrillary tangles (NFTs). Additionally, WWOX possesses proapoptotic properties, and its loss-of-function has been shown to result in the activation of a protein aggregation cascade [30]. *ETNK1* encodes an ethanolamine-specific kinase involved in phosphatidylethanolamine biosynthesis, which is crucial for the folding and activity of several membrane proteins, initiation of autophagy, and maintaining optimal mitochondrial respiratory activity and ubiquinone function [19, 47]. The increased methylation of two CpGs in the promoter region of *ETNK1* suggests a dysregulation/repression of *ETNK1,* potentially leading to low phosphatidylethanolamine levels, resulting in protein misfolding and aggregation due to abnormal protein degradation (impaired autophagy) that is characteristic of these diseases. *FAM8A1* encodes a protein that is associated with ubiquitin dependent endoplasmic reticulum-associated degradation of proteins with roles in AD pathogenesis, a differentially methylated CpG mapping to *FAM8A1* was also the most significantly associated with AD in a previous study [61]. Additionally, *DFNA5,* the hub gene identified in the WGCNA module commonly associated with the three parkinsonian disorders, is a pyroptotic gene reported to induce programmed cell death through mitochondria and MAPK related pathways [60] and mediates mitochondrial damage in axons and neurodegeneration [44]. Differential methylation of these loci commonly identified in the parkinsonian diseases suggests commonalities in terms of pathways related to autophagy, mitophagy, apoptosis, and protein degradation pathways in all these diseases.

Among the CpGs identified in the individual comparisons of each disease group with controls, hypomethylation in the promoter region at cg15644686 mapping to *BCL7B* was observed in MSA, with concordant transcriptional upregulation being reported in the cerebellar white matter. BCL7B (BAF chromatin remodelling complex subunit BCL7B) is a negative regulator of Wnt signalling and a pro-apoptotic factor, and a deficiency in BCL7B reportedly enhances oligodendrogenesis [27, 59, 68]. This may suggest that increased levels of BCL7B in MSA might hinder oligodendrogenesis. This, in conjunction with the gliosis and demyelination observed in MSA, could exacerbate disease pathology, as indicated by the inverse correlation between methylation levels and disease duration and the higher mean number of GCIs in MSA as we observed in our study. The CpG cg01380065 showed hypermethylation in PD compared to controls. This CpG maps to *UBE2F*, which has been shown to be involved in neddylation, which is a post-translational modification essential for regulating the clearance of misfolded proteins [24]. The CpG cg25358066, found to be hypermethylated in with the strongest effect in PSP, mapped to *D2HGDH* (D-2-hydroxyglutarate dehydrogenase), a mitochondrial enzyme belonging to the FAD-binding oxidoreductase/transferase type 4 family and an overexpression of D2HGDH has been demonstrated to inhibit ferroptosis [65].

Analysis of the differential methylation signatures with potential to discriminate between the diseases performed by comparing the disease groups against each other revealed that the extent of disease specificity in terms of differential methylation between these diseases in the white matter is limited. Although a few differentially methylated CpGs that showed opposite direction of effect in one disease compared to the other were identified such as *FMO6P* and *OR51A7* in MSA compared to PD, in most cases, dysregulation was still observed with the CpG in all three disease groups. Notable hub genes specifically identified in modules correlating with α-synucleinopathies included *C1orf70,* which has been implicated to play a role in spinocerebellar ataxia, and *SCARF2,* which maps to the 22q11 deletion region previously associated with increased PD risk, suggesting similar roles of these genes in MSA pathogenesis [38].

Co-methylation modules enriched for oligodendrocytic genes included some modules commonly associated with all three parkinsonian diseases, and some that were associated with synucleinopathies only in addition to some disease-specific modules. The oligodendrocyte enriched module darkgrey, significantly positively associated with PD, PSP and to a certain extent MSA, showed functional enrichment of processes such as RNA interference (RNAi), signal transduction, ER stress, mitochondrial translation, and endosomal transport suggesting a common involvement of these molecular pathways in all three parkinsonian diseases. Mechanisms relating to RNAi have already been reported for several neurodegenerative diseases including in PD, where mutations in genes encoding proteins responsible for cellular import of dsRNA in a *C*. *elegans* model reportedly protect dopamine neurons from degeneration [20]. Moreover, therapeutic models of RNAi are being extensively studied, with short hairpin (sh) RNA being the most commonly used construct in animal models of HD, AD, and PD [21]. Intra- or inter-cellular signalling mechanisms have also been described to be involved in the pathogenesis of neurodegenerative diseases with effectors and/or components of the signal transduction pathways playing important roles in progression, and possibly in the initiation, of these diseases [64]. Mechanisms relating to ER stress, mitochondrial functions and endosomal transport have been extensively reported in neurodegenerative diseases including PD, with PARK17 playing a role in the retrotransfer of proteins from endosomes in the pre-lysosomal compartment network to the trans-Golgi network, and PARK9 and ATP13A2 coding for endo-/lysosomal related proteins, HTRA2 (PARK13) being crucial to maintaining normal mitochondrial function and ERS-coupled apoptotic cell death being implicated in neurodegeneration [13, 58].

The oligodendrocyte-enriched white module, which was significantly associated commonly in the synucleinopathies (MSA and PD), showed enrichment of processes relating to the Wnt signalling pathway. Wnt signalling pathway has previously been shown to play an important role in PD pathogenesis, with dysfunction in PARKIN, leading to the accumulation of β-catenin and resulting in the upregulation of canonical Wnt signalling. Interestingly, *PARKN* was a hub gene in the M3 functional module of the white module and also contained our MSA top hit *BCL7B,* which has been shown to play a role in the Wnt signalling pathway by negatively regulating the expression of Wnt signalling components CTNNB1 and HMGA1 [59]. Put together, these findings indicate a dysregulation in Wnt signalling pathways to be common in both MSA and PD even in the white matter. Moreover, white matter damage has been found to precede grey matter atrophy in PD [1, 15]. Wnt signalling pathways play important roles in oligodendrogenesis, oligodendrocyte differentiation, and myelination, and DNA methylation alterations dysregulating the Wnt signalling pathway might be one of the factors responsible for preventing remyelination through the mobilisation of OPCs following death of oligodendrocytes or myelin damage due to disease [23, 52].

Modules enriched for oligodendrocytes significantly associated with PSP included the darkred and steelblue modules, both of which also showed inverse associations with synucleinopathies. Both these modules showed enrichment in processes related to RNA splicing, mRNA and peptide metabolic processes; the darkred module was also enriched for histone methylation and positive regulation of tumour necrosis factor (TNF)-mediated signalling pathway. The inverse correlation observed between the synucleinopathies and taupathy could be attributed to differences in these molecular processes or different molecular players within these processes driving the pathogenesis. Additionally, the darkred module also showed enrichment for the Wnt signalling pathway, which has also been reported in a previous DNA methylation study conducted in PSP forebrains [66], suggesting that DNA methylation alterations within this pathway is a common factor in white matter tissues across MSA, PD, and PSP. The MSA associated midnightblue module, in addition to being enriched for the commonly identified Wnt signalling and apoptotic processes, also showed enrichment in ER pathways such as COPII-coated vesicle budding and cargo loading into COPII-vesicle, protein dephosphorylation, and cytokine-mediated signalling pathway suggesting additional roles of these pathways in MSA pathogenesis. Microglial activation has been shown to parallel system degeneration in MSA, where these microglia express numerous cytokines and other inflammation-associated molecules [29]. However, activated microglia have also been observed in PD, and glial cytokine-mediated, neuroimmunologic processes underlie the progression of several neurodegenerative diseases indicating that dysregulation in these processes is not exclusive to MSA [40].

As any other DNA methylomic study, our study also has several limitations. Although the analysis of frontal lobe white matter DNA methylation profiles revealed several commonalities in MSA, PD, and PSP, brain regions and tissue-types primarily affected in the three diseases vary. DNA methylation differences in the primary affected region in each disease, such as substantia nigra/basal ganglia in PD, striatum, substantia nigra and cerebellum in MSA, and subcortical and cortical regions in PSP, could not be captured in this study as we focused on the frontal lobe white matter. However, we chose the frontal lobe, as it shows moderate to high pathology at the end-stage in all three diseases, to be able to perform a cross-disease comparison. Therefore, further studies examining the extent of dysregulation of the identified DNA methylation alterations in other brain regions might provide additional insights into disease-specific patterns. Moreover, as our study uses post-mortem brain tissues, we cannot identify early changes in DNA methylation, and cannot distinguish the causative DNA methylation alterations from the reactive changes. Our modest sample sizes per group only made it possible to identify differential DNA methylation alterations only at nominal significance. However, we compensated this limitation by employing more powerful system biology approaches such as co-methylation networks to further complement and strengthen our findings.

In conclusion, our study provides the first evidence of DNA methylation alterations in the white matter of three different parkinsonian disorders that point to common DNA methylation alterations and shared pathogenic mechanisms in the three diseases. Our study reveals the overall presence of more similarities than differences in MSA, PD, and PSP frontal lobe white matter in terms of DNA methylation architecture, with differences between diseases primarily lying in the effect sizes of the alterations. While this study identifies shared mechanisms that provide valuable insights into the DNA methylation perturbations in parkinsonian disorders, further studies involving larger sample numbers and multiple regions and tissue/cell types of the brain are warranted for the identification of disease-specific methylation perturbations in the diseases. Nevertheless, our study identifies several candidate loci that display shared DNA methylation dysregulation in the frontal lobe white matter in all three Parkinsonian diseases that can be further explored as potential therapeutic targets and highlights common pathogenic mechanisms between the diseases, which are indicative of converging molecular pathways that contribute to neurodegeneration in MSA, PD, and PSP.

## Acknowledgements

The authors would like to thank UCL Genomics centre for processing the EPIC arrays for the frontal lobe white matter dataset. The authors would also like to acknowledge the Queen Square Brain Bank (London, UK) for providing brain tissues from disease cases and controls. The Queen Square Brain Bank is supported by the Reta Lila Weston Institute of Neurological Studies, UCL Queen Square Institute of Neurology. MM and NR are supported by a grant from the Multiple System Atrophy Trust awarded to CB. KF is supported by the Medical Research Council (MR/N013867/1). TL is supported by an Alzheimer’s Research UK Senior Fellowship. TW is supported by the Reta Lila Weston Trust and the MRC (N013255/1). CB is supported by Alzheimer’s Research UK (ARUK-RF2019B-005) and the Multiple System Atrophy Trust.

## Data Availability Statement

Raw methylation data for the MSA prefrontal cortex, LBD, and PSP prefrontal lobe datasets are available in NCBI GEO database (https://www.ncbi.nlm.nih.gov/geo), and can be accessed via accession numbers GSE143157, GSE203332, GSE197305, and GSE75704. Additional data is available in supplementary materials and from the corresponding author upon reasonable request.

