## Supplementary figures for "DNA methylation patterns in the frontal lobe white matter of multiple system atrophy, Parkinson’s disease, and progressive supranuclear palsy: A cross-comparative investigation"

United Kingdom

### Supplementary Figures

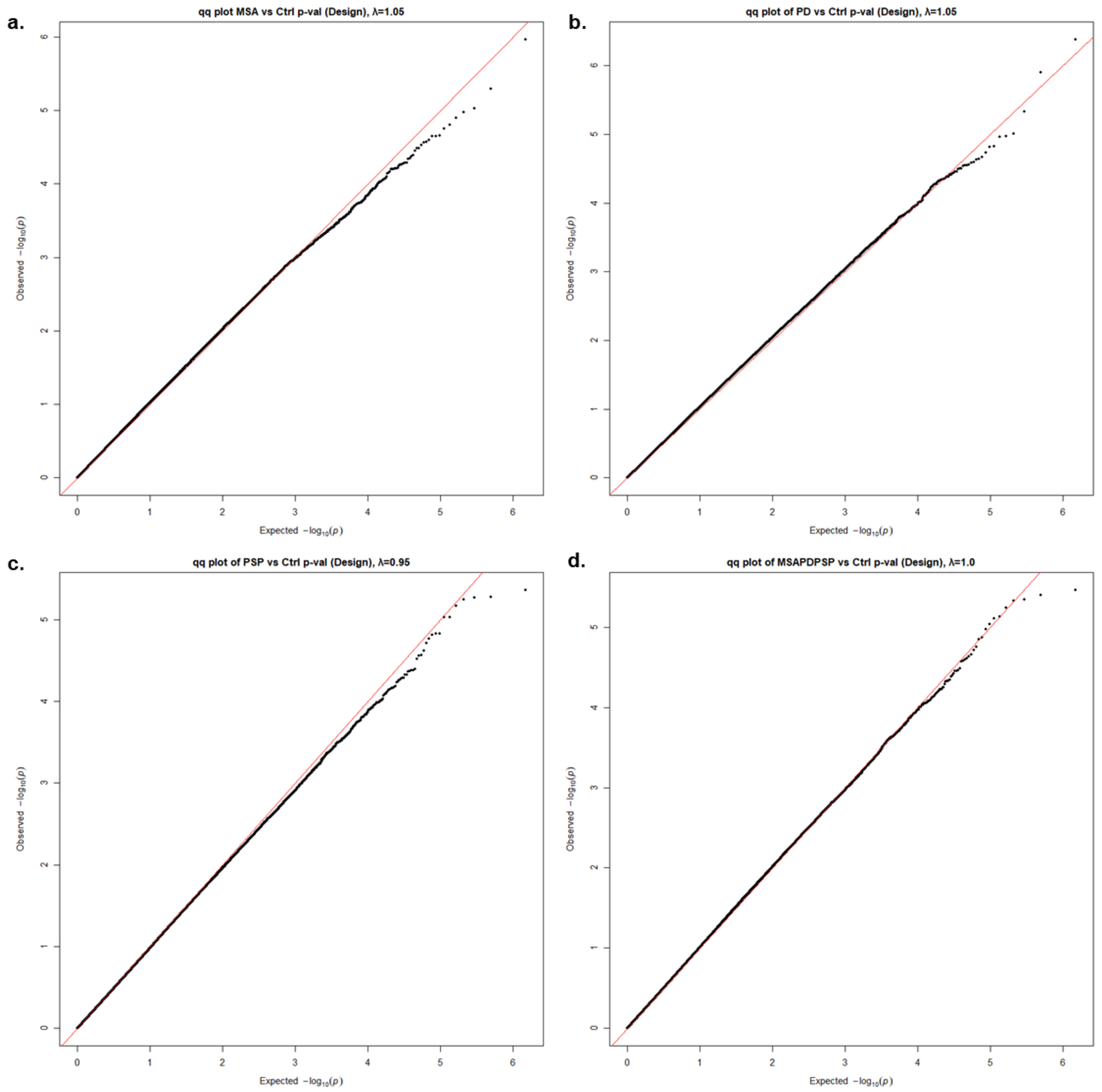


**Figure S1:** Quantile-quantile (Q-Q) plots for the disease group specific case-control EWAS (a) Q-Q plot for MSA shows an estimated inflation factor (λ) of 1.05, (b) Q-Q plot for PD shows an estimated inflation factor (λ) of 1.05, (c) Q-Q plot for PSP shows an estimated inflation factor (λ) of 0.95, (d) Q-Q plot for all disease groups shows an estimated inflation factor (λ) of 1.0. CTRL – controls, MSA – multiple system atrophy, PD – Parkinson's disease, PSP – progressive supranuclear palsy.

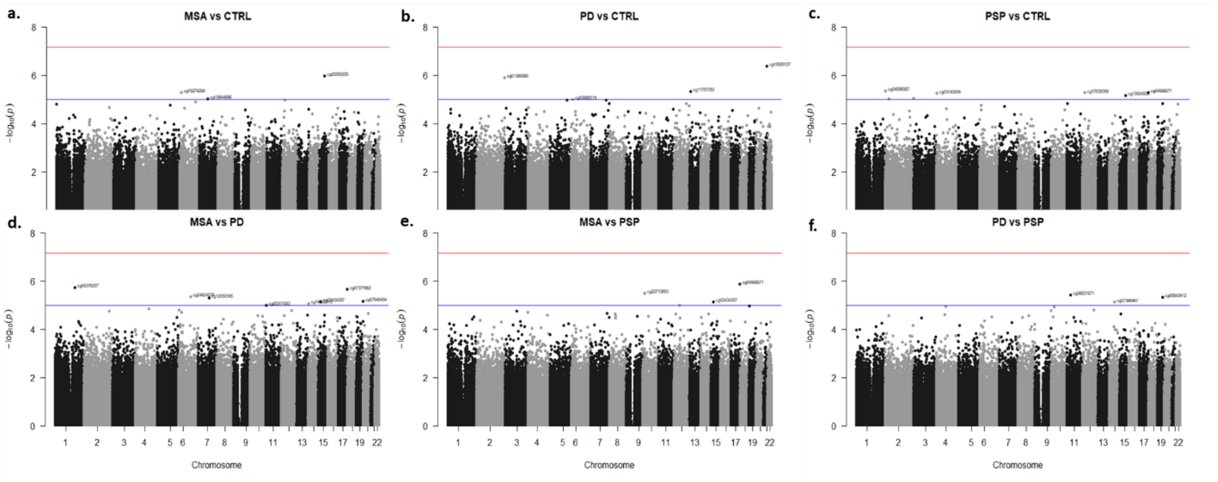


**Figure S2:** Manhattan plots showing the p-value distributions of the differentially methylated probes for the different comparisons, (a.) MSA vs CTRL, (b.) PD vs CTRL, (c.) PSP vs CTRL, (d.) MSA vs PD, (e.) MSA vs PSP, and (f.) PD vs PSP. The red line indicates genome-wide significance threshold based on Bonferroni-corrected p-values ($p = 6.8 \times 10^{-8}$), and the blue line indicates a less stringent suggestive significance threshold of $p = 1 \times 10^{-5}$. CTRL – controls, MSA – multiple system atrophy, PD – Parkinson's disease, PSP – progressive supranuclear palsy.

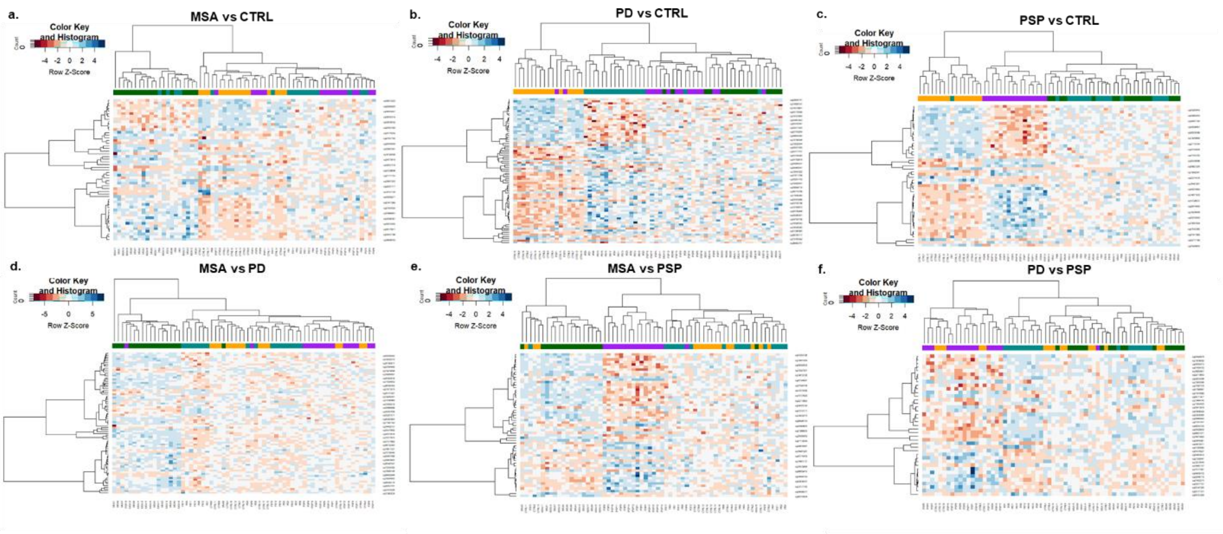


**Figure S3:** Heatmap of the topmost significant differentially methylated loci ($p < 0.0001$) identified in (a.) MSA compared to controls, (b.) PD compared to controls, (c.) PSP compared to controls, (d.) MSA compared to PD, (e.) MSA compared to PSP, and (f.) PD compared to PSP. The rows represent CpGs, columns represent samples, and the colours represent the direction as well as the magnitude of effect (adjusted β values) in all the samples.

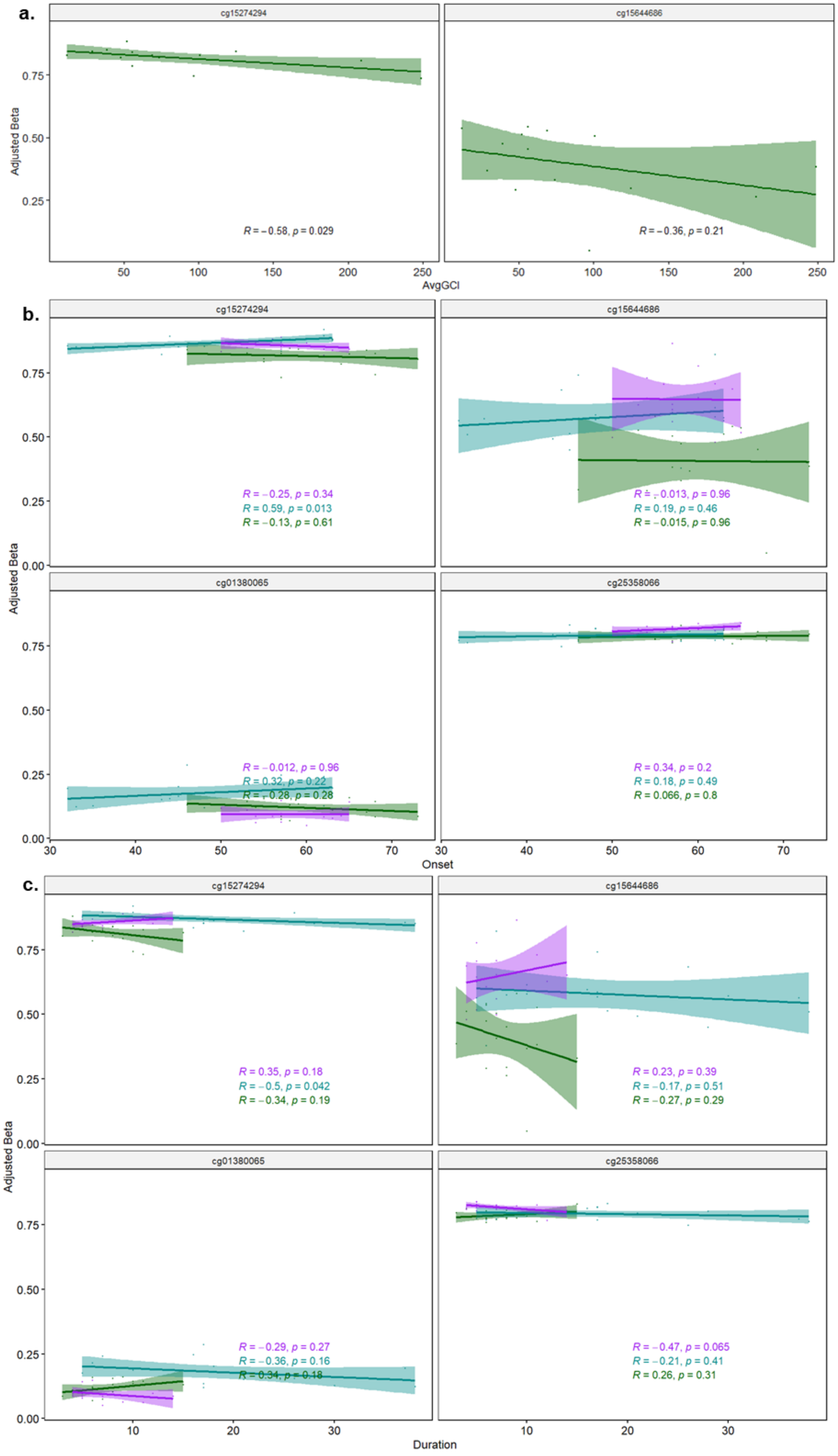

a.
b.
c.
cg15274294
cg15644686
cg01380065
cg25358066
Adjusted Beta
AvgGCI
Onset
Duration
R = −0.58, p = 0.029
R = −0.36, p = 0.21
R = −0.25, p = 0.34
R = 0.59, p = 0.013
R = −0.13, p = 0.61
R = −0.013, p = 0.96
R = 0.19, p = 0.46
R = −0.015, p = 0.96
R = −0.012, p = 0.96
R = 0.32, p = 0.22
R = −0.28, p = 0.28
R = 0.34, p = 0.2
R = 0.18, p = 0.49
R = 0.066, p = 0.8
R = 0.35, p = 0.18
R = −0.5, p = 0.042
R = −0.34, p = 0.19
R = 0.23, p = 0.39
R = −0.17, p = 0.51
R = −0.27, p = 0.29
R = −0.29, p = 0.27
R = −0.36, p = 0.16
R = 0.34, p = 0.18
R = −0.47, p = 0.065
R = −0.21, p = 0.41
R = 0.26, p = 0.31

**Figure S4:** Correlation between differential methylation levels and disease associated traits for the DMPs cg15274294 (Chr 6 – IGR), cg15644686 (*BCL7B*), cg01380065 (*UBE2F*), and cg25358066 (*D2HGDH*) in the different disease comparisons. Scatter plot and trend line (Pearson's correlation) showing correlation between methylation levels and (a) average GCI, (b) disease onset, and (c) disease duration. MSA– multiple system atrophy (mixed subtype), PD – Parkinson's disease, PSP – progressive supranuclear palsy, AvgGCI – average number of glial cytoplasmic inclusions.

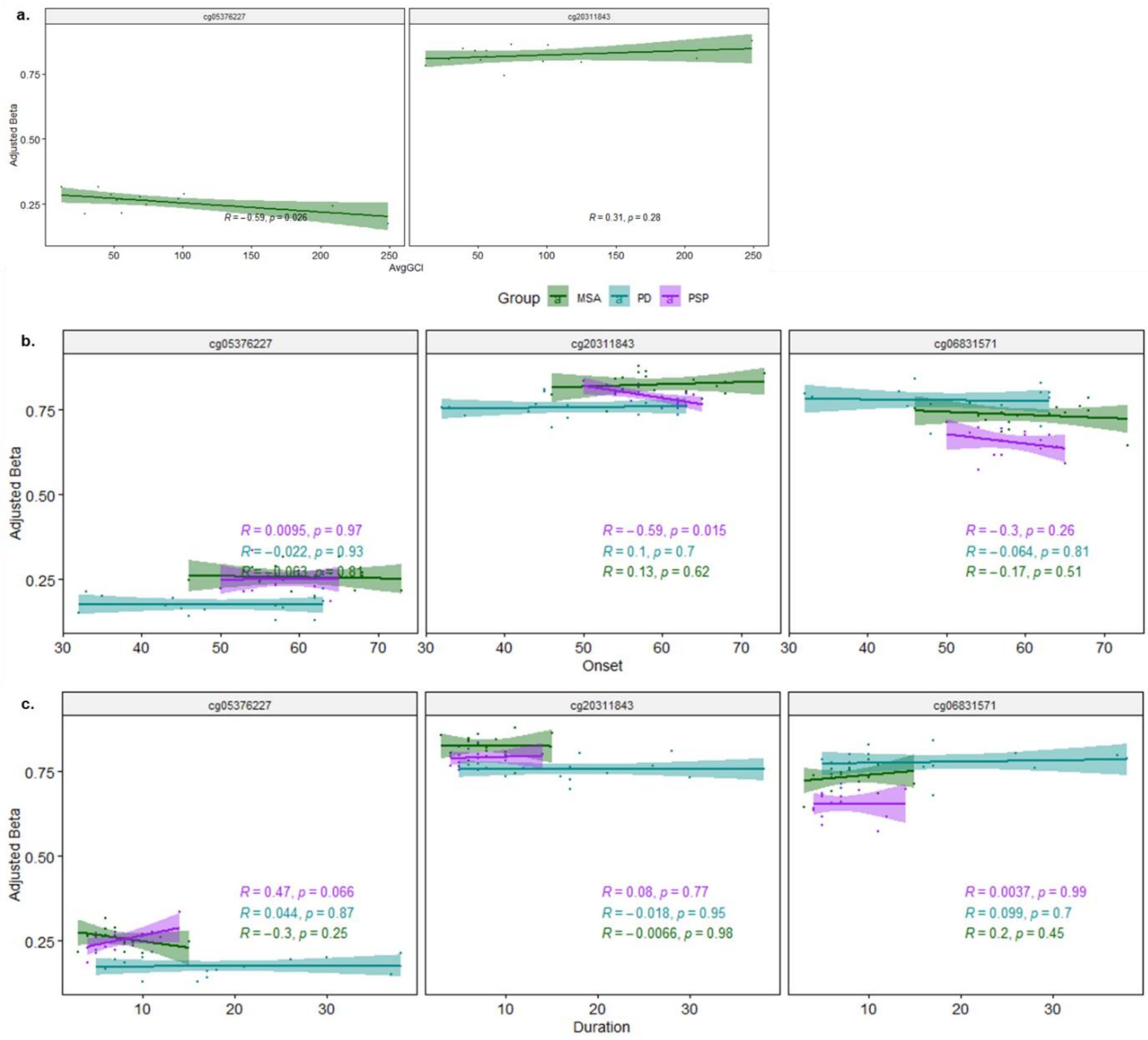


**Figure S5:** Correlation between differential methylation levels and disease associated traits for the DMPs cg05376227 (*FMO6P*), cg20311843 (*OR51A7*) in MSA vs PD and cg06831571 (Chr11 – IGR) in PD vs PSP; scatter plot and trend line (Pearson's correlation) showing correlation between methylation levels and (a) average GCI, (b) disease onset, and (c) disease duration. MSA– multiple system atrophy (mixed subtype), PD – Parkinson's disease, PSP – progressive supranuclear palsy, AvgGCI – average number of glial cytoplasmic inclusions.

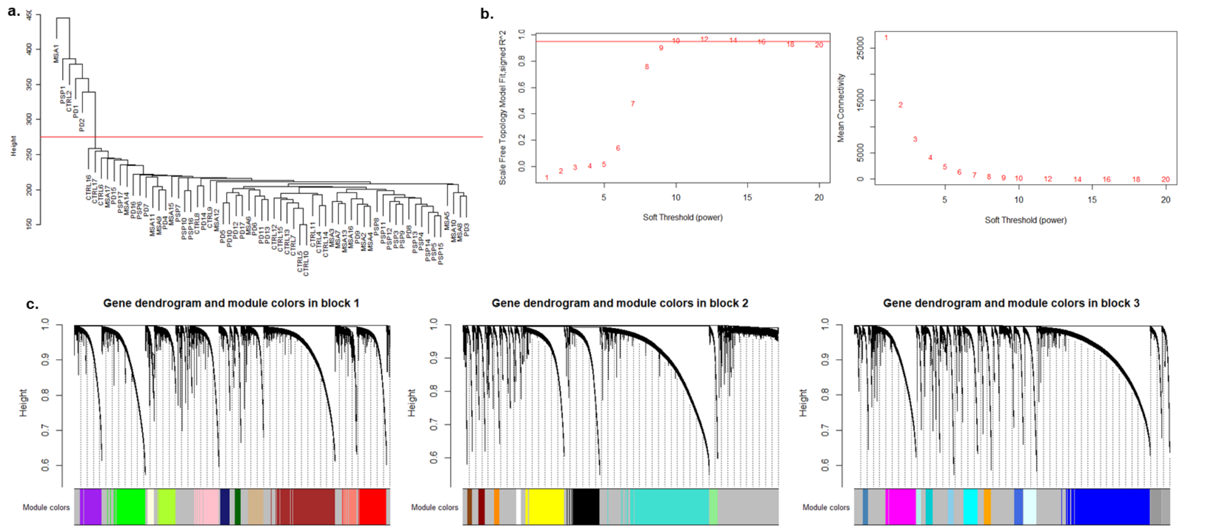


**Figure S6:** Cluster trees and scale-free topology criteria for the co-methylation network: (a) Sample clustering to detect outliers, (b) scale-free topology (SFT) plot for choosing the soft-thresholding power β for the signed weighted correlation network; left: SFT index $R^2$ (y-axis) as a function of different powers β (x-axis), right: mean connectivity (y-axis) as a function of the power β (x-axis), (c) hierarchical clustering dendrogram generated using blockwise module detection.

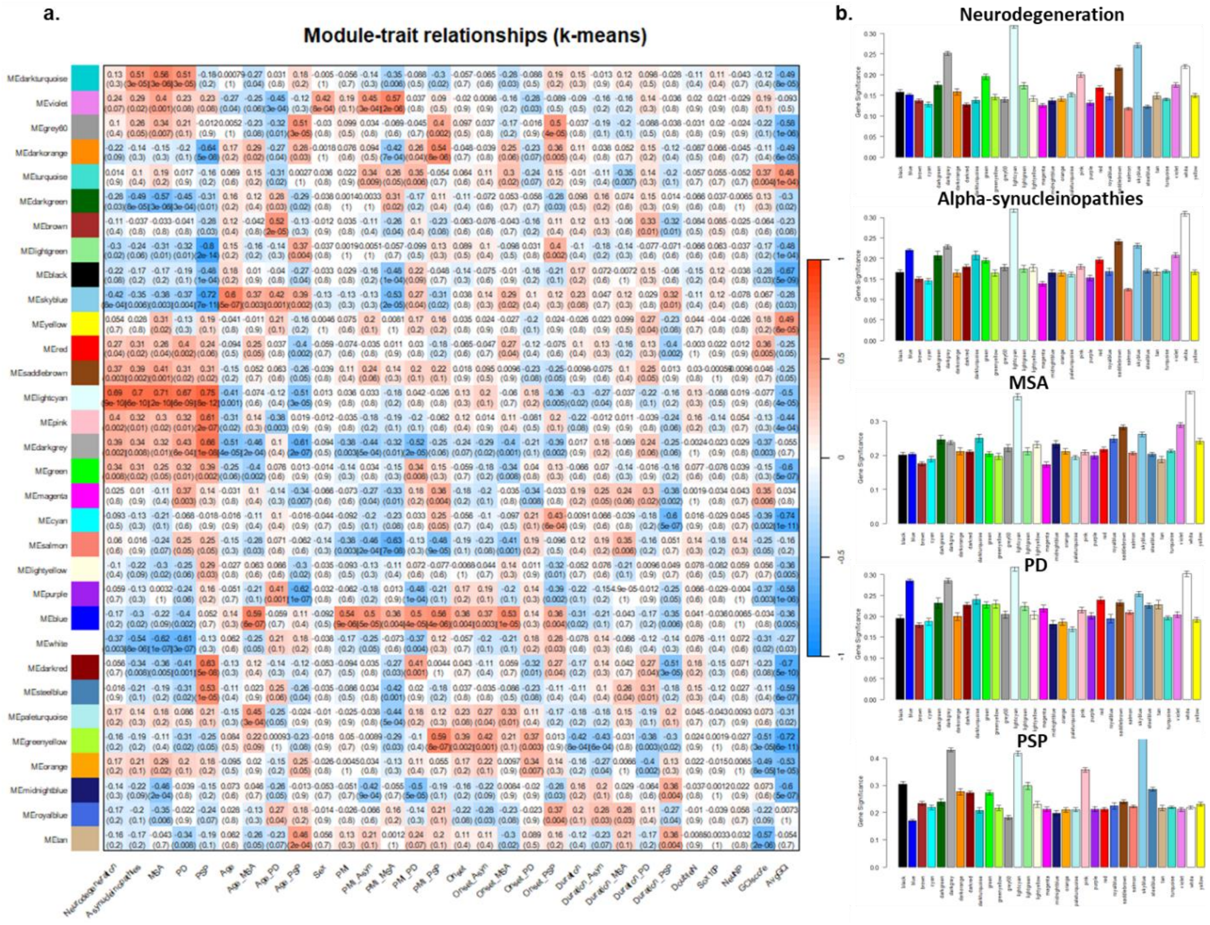


**Figure S7:** Module-trait correlations and gene significance. (a) heatmap showing the module-trait correlations and p values for all disease associated clinical/pathological traits, (b) gene significance of the different modules in all three disease groups (neurodegeneration), in MSA and PD (α-synucleinopathies), and in the individual disease groups – MSA, PD, and PSP. MSA– multiple system atrophy, PD – Parkinson’s disease, PSP – progressive supranuclear palsy.

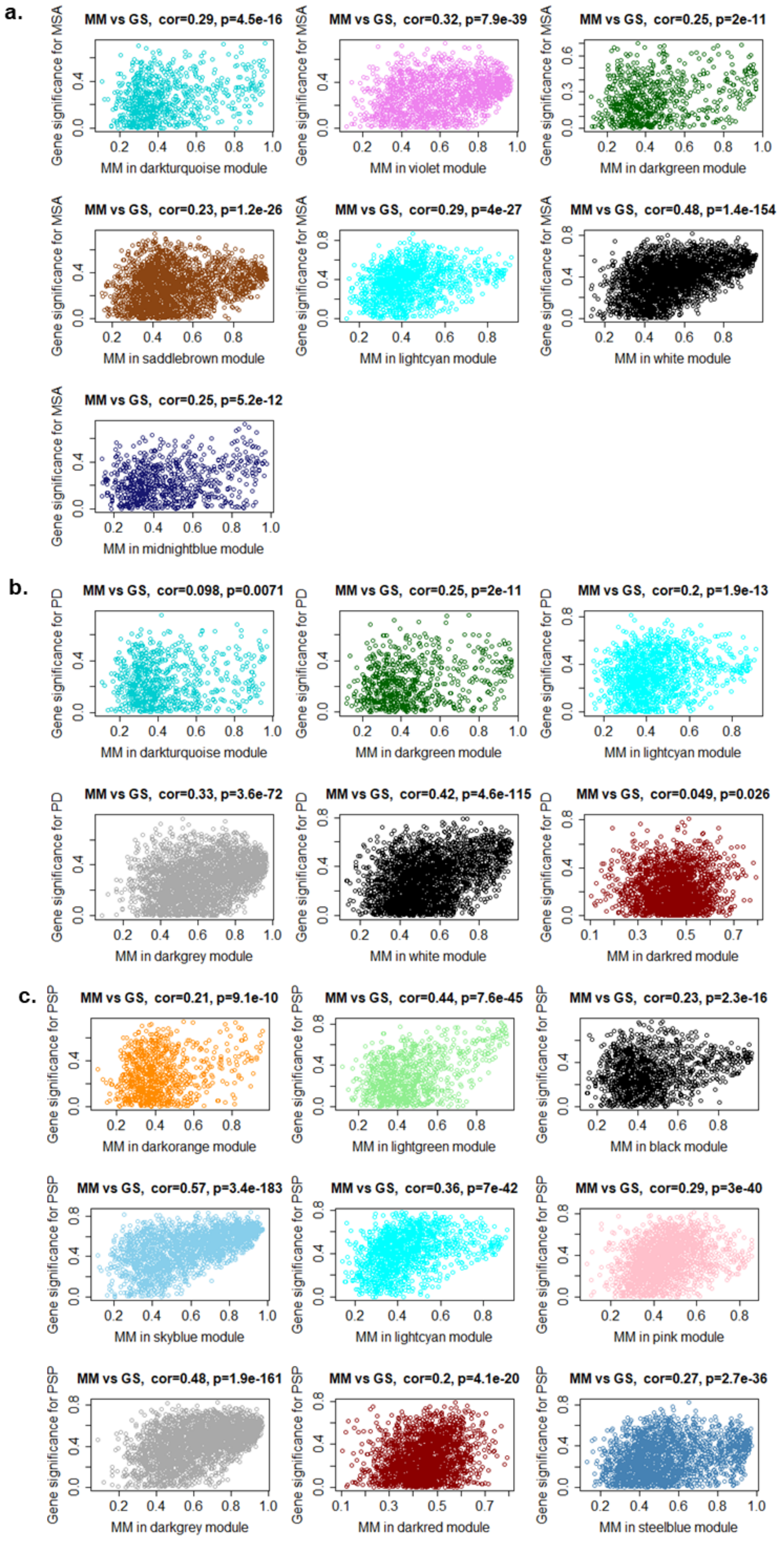


**Figure S8:** Correlation between gene significance and module membership (kME) for modules significantly associated (adj.p≤0.001) with (a) MSA, (b) PD, and (c) PSP. MSA– multiple system atrophy,

PD – Parkinson’s disease, PSP – progressive supranuclear palsy, GS – gene significance, MM – module membership.
